# Temporally precise movement-based predictions in the mouse auditory cortex

**DOI:** 10.1101/2021.12.13.472457

**Authors:** Nicholas J. Audette, WenXi Zhou, David M. Schneider

## Abstract

Many of the sensations experienced by an organism are caused by their own actions, and accurately anticipating both the sensory features and timing of self-generated stimuli is crucial to a variety of behaviors. In the auditory cortex, neural responses to self-generated sounds exhibit frequency-specific suppression, suggesting that movement-based predictions may be implemented early in sensory processing. Yet it remains unknown whether this modulation results from a behaviorally specific and temporally precise prediction, nor is it known whether corresponding expectation signals are present locally in the auditory cortex. To address these questions, we trained mice to expect the precisely timed acoustic outcome of a forelimb movement using a closed-loop sound-generating lever. Dense neuronal recordings in the auditory cortex revealed suppression of responses to self-generated sounds that was specific to the expected acoustic features, specific to a precise time within the movement, and specific to the movement that was coupled to sound during training. Predictive suppression was concentrated in L2/3 and L5, where deviations from expectation also recruited a population of prediction-error neurons that was otherwise unresponsive. Recording in the absence of sound revealed abundant movement signals in deep layers that were biased toward neurons tuned to the expected sound, as well as temporal expectation signals that were present throughout the cortex and peaked at the time of expected auditory feedback. Together, these findings reveal that predictive processing in the mouse auditory cortex is consistent with a learned internal model linking a specific action to its temporally precise acoustic outcome, while identifying distinct populations of neurons that anticipate expected stimuli and differentially process expected versus unexpected outcomes.

## Introduction

As animals interact with the world, many of the sensations they encounter are caused by their own actions. Accurately anticipating both the sensory features and timing of these self-generated stimuli is crucial to a variety of behaviors, such as using auditory input to guide ongoing vocalizations, or determining if footsteps are self-generated or belong to a predator. The brain’s ability to predict the sensory consequences of actions is thought to involve the integration of internally generated motor signals with incoming sensory information using learned internal models ^1–10^. This computation, referred to as predictive processing, is hypothesized to improve sensory processing by diminishing neural responses to expected stimuli while enhancing neural responses to new or unexpected stimuli, and provides a viable substrate for sensory-guided motor learning ^3,11–14^.

A growing body of experimental evidence supports the theory that predictive processing is implemented in the sensory cortex, where neural activity is strongly influenced by behavior ^5,7,21–23,8,9,15–20^. Sensory cortical regions receive dense input from motor-related regions that exert a strong influence over spontaneous and stimulus-evoked activity during movement ^2,4,24–27^. For example, in the mouse auditory cortex, sound-evoked responses are suppressed during movement compared to rest, mediated in part through long-range inputs from the motor cortex onto local inhibitory cells ^4,22^. Importantly, movement-related modulation in the cortex can reflect specific expectations. When a movement reliably produces a sound, suppression in the auditory cortex becomes stronger for self-generated sounds that match the expected frequency, and neurons respond more strongly when a movement produces an unexpected frequency ^28–32^. Neurons in the mouse visual cortex also respond more strongly when a behavior is accompanied by unexpected visual consequences, further supporting the idea that prediction may be a fundamental component of cortical sensory processing ^24,33–35^.

However, many observations that resemble predictive processing may be interpretable through non-predictive mechanisms, such as behavioral-state-dependent gain control of visual cortex tuning curves^36^. Stimulus-specific modulation can also result from short-term reactions to recent stimulus patterns or from longer time-course expectations related to an animal’s general behavioral state rather than a specific movement ^4,37^. If the sensory cortex implements specific movement-based predictions, several important features should be present: the confinement of predictive suppression to a specific behavior; the capacity to detect changes in stimulus timing with within-movement precision; and the local presence of predictive signals that encode the expected timing and identity of a sensory outcome. Establishing these principles in a single experimental paradigm is critical to determining whether motor-sensory predictions are implemented in the sensory cortex but has been challenging because it requires gaining experimental control over the precise sensory consequences of an isolated and stereotyped movement.

To test whether these features of predictive processing are present in the auditory cortex, we developed a simple behavioral paradigm in which mice experience consistent auditory feedback while performing a forelimb movement that is newly learned, intermittent rather than continuous, and progresses over an ethologically relevant timescale of hundreds of milliseconds^38^. Dense neuronal recordings in the auditory cortex of trained mice revealed learned suppression of expected, self-generated sounds that was frequency-specific, temporally precise, and restricted to the sound-associated movement. By recording in the absence of sound, we also identified correspondingly precise movement signals that reflected a learned expectation for the identity and timing of the expected sound. These signals were distributed across functional groups of neurons with distinct laminar patterns, including predictive suppression and prediction error neurons concentrated in L2/3 and L5, movement-responsive neurons concentrated in L6, and temporally precise expectation neurons distributed across cortical layers. Together, these findings illustrate layer-specific predictive processing in the auditory cortex that is consistent with a learned internal model linking a specific action to its temporally precise acoustic outcome.

## Results

### Frequency-specific suppression of expected self-generated sounds

We designed a behavioral paradigm in which head-fixed mice learn to push a lever with their forelimb while receiving closed-loop auditory feedback (Fig 1A). All movements were mouse initiated, with lever pushes that crossed a fixed threshold (~15 mm) resulting in a small water reward when the lever was returned to a resting position (Fig 1B). A brief pure tone was delivered at a fixed position early in the outbound phase of each movement creating an experimentally controlled and consistent sensory outcome for each lever push. Movements that closely followed a previous lever push (< 200ms inter-trial interval) generated a sound but did not result in a reward, encouraging a slower rate of trial initiation. No other shaping criteria were applied. Mice rapidly learned to make prolonged lever movements that reached the target threshold, lasting an average of 365ms with a median inter-trial interval of 3.5s (Fig 1C). Across training, mice made thousands of lever pushes (referred to as trials) and heard thousands of self-generated sounds at a specific position of their movement.

**Figure 1.**
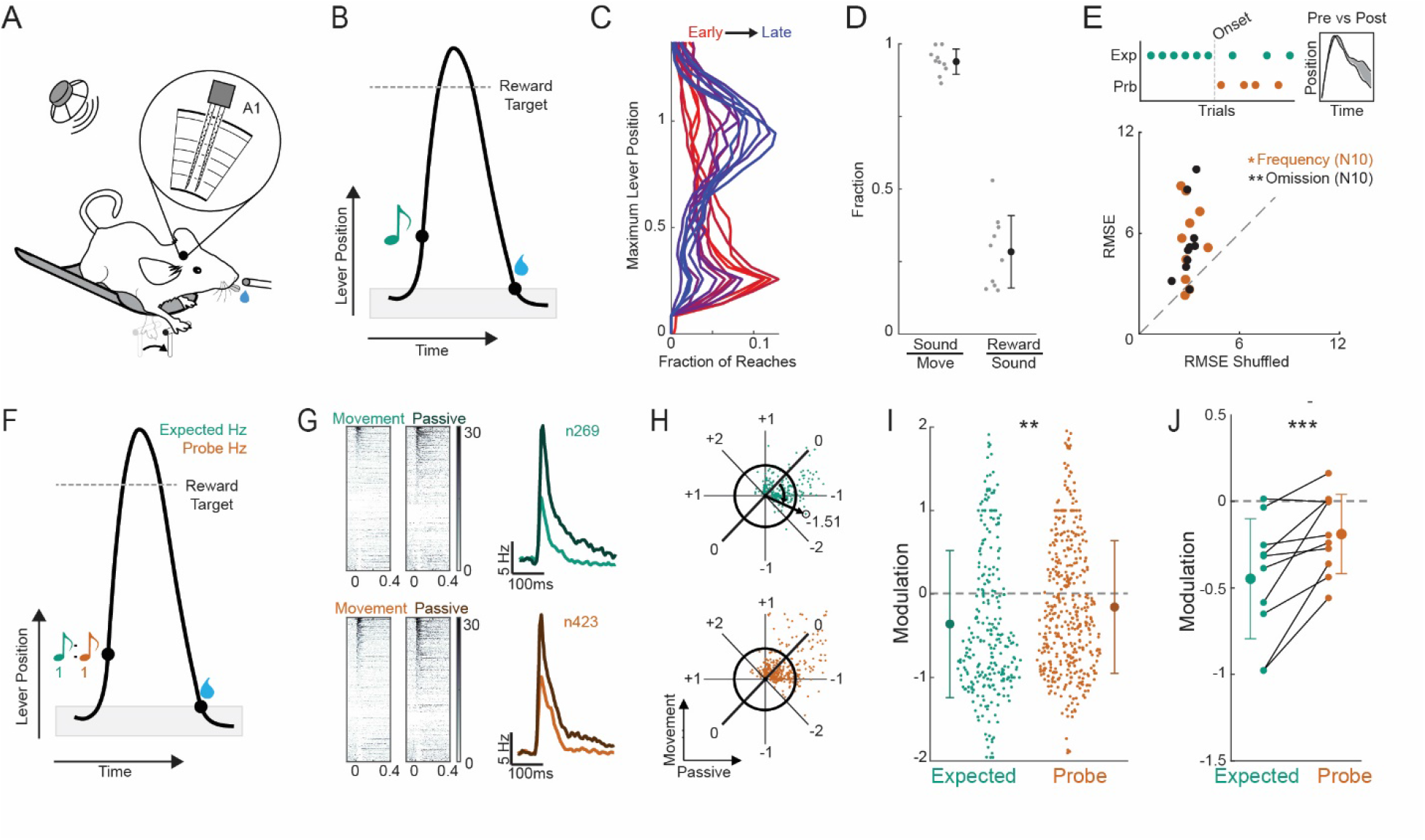
Frequency-specific suppression of self-generated sounds in the mouse auditory cortex. (A) Schematic of multi-array recordings during head-fixed lever press paradigm where mice hear closed-loop auditory feedback while making rewarded forelimb movements. (B) Schematic of stimulus and reward timing for each lever movement. Grey box indicates lever home position. (C) Histogram of reach distances normalized to the position of the reward threshold across training sessions (red, early; blue, late) for an example mouse. (D) Movement, sound, and reward coupling measured as the fraction of movements that generated sounds (crossed the sound threshold), and the fraction of sound-generating movements that generated rewards (obeyed inter-trial-interval and reached reward threshold)(N=10 mice). (E) Schematic (top) of introduction of deviant auditory feedback (probe) during lever press task, and measurement of lever trajectory RMSE difference between the 30 trials prior to the onset of deviants and after the onset of deviants, regardless of trial type. Quantification (bottom) of lever trajectory RMSE differences before and after the onset of frequency deviant (orange) or omission (black) trials, compared to the average RMSE difference between groups comprised of the same trials, but assigned random identities (mean of 1000 permutations). (F) Schematic of frequency-probe session where mice hear either the expected frequency or a probe frequency one octave away on 50% of trials. (G) Heat maps (left) of all neural responses (n=928, N=10) to the lever-associated sound (top, cyan) and frequency probe sound (bottom, orange) in the active condition (light, left) and passive condition (dark, right). PSTHs (right) of neural response to lever-associated sound (top, cyan) and frequency probe sound (bottom, orange) in the active condition (light) or passive condition (dark) averaged across all recorded neurons responsive to that sound in either the active or passive condition. (H) Scatter of individual neural responses to either the expected (cyan) or probe (orange) tone in the active condition vs passive condition. Schematic of radial modulation index calculation is overlayed, with a highlighted example neuron and its corresponding modulation value. (I) Modulation index of individual neurons comparing active and passive responses to the expected and probe frequencies. (J) Connected points display the average neural modulation of expected and probe frequencies for an individual animal. *p<0.05 **p < 0.005 ***p<.0005

Our task was designed so that mice would learn an expectation for the timing and identity of a self-generated tone, without relying on the tone to complete the task or signal reward. Consistent with this design, mice heard a sound on almost every movement of the lever, while rewards were delivered on a smaller subset of trials on which mice crossed both the sound and the reward threshold (Fig 1D). To directly test whether mice developed an expectation for the self-generated tone, we measured lever trajectories and licking behavior when the lever-associated tone was altered (Fig S1A-C) or omitted (Fig S1D-F) on a subset of trials. We observed no consistent change in lever movement or licking behavior on trials with deviant acoustic feedback compared to interleaved control trials (FIG S1A-F). However, when measuring longitudinally we observed a transient change in the overall shape of lever trajectories upon the initiation of deviant auditory feedback, measured as the root-mean-square error (RMSE) difference between the average movement lever trajectory for trials just before and just after the start of unexpected outcomes (Fig 1E). Since subtle behavioral differences were visible before and after the introduction of deviants, but not across interleaved trial types, we conclude that mice do not rely on the tone to perform lever reaches but do possess an expectation for the presence and identity of the expected sound.

We next asked how this learned expectation influences sensory processing in the auditory cortex by comparing neural responses to self-generated sounds with expected and unexpected frequencies. Following behavioral training, a 128-channel electrode array was lowered into the auditory cortex, oriented perpendicular to the cortical surface (Fig 1A)^39^. The lever was introduced as normal, and mice began making volitional lever presses while experiencing expected auditory feedback. After collecting auditory cortical responses during lever movements that produced expected self-generated sounds, we unexpectedly introduced frequency probe trials, where auditory feedback occurred at the correct position but with tones shifted one octave on 50% of randomly interleaved trials (Fig 1F). At the end of the recording session, the lever was removed, and mice were presented with the same auditory stimuli in a passive listening context to provide baseline sound response properties of all recorded neurons. In total, we recorded passive and self-generated sound responses from 928 well-isolated, non-fast-spiking units across 10 animals.

Across the entire neural population, the average strength of neural responses to the lever-associated frequency and probe frequency heard in the passive condition were comparable (Fig 1G, ranksum, p = 0.79). When the same tones were generated by the mouse’s lever movement, neural responses to the expected and probe frequencies were both suppressed, but suppression was strongest for the expected self-generated frequency (Fig 1G). To specifically quantify the activity of neurons that were sound responsive, subsequent analyses included only neurons that responded to a given frequency in at least one of the active or passive conditions. Peri-stimulus time histograms (PSTHs) of these sound-responsive populations revealed that neural responses to the expected frequency were diminished by 62% compared to passive listening (Fig 1G, 5.2 Hz vs 13.6 Hz), while neural responses to the unexpected frequency displayed weaker suppression of 41% (Fig 1G, 8.5 Hz vs 14.2 Hz). These coarse population-level analyses indicate the presence of at least two forms of movement-related suppression within the cortex: a generic suppression that applies to all sounds and a predictive component that is specific to the expected sound frequency^28^.

To determine the degree to which movement-related modulation of sound-evoked responses varied across the neural population, we calculated a radial modulation index for each neuron and for each of the expected and unexpected frequencies (Fig 1H). Negative modulation indices indicate neurons with weaker responses when the sound is self-generated compared to the same sound heard passively, whereas positive modulation indices indicate neurons that respond more strongly to self-generated sounds. Consistent with population-level PSTHs, most neurons were suppressed for both frequencies (Fig 1I). However, average modulation was more negative for the expected frequency (−0.36) than for the probe frequency (−0.17) across all neurons (Fig 3I), and in 9 out of 10 animals (Fig 1J). Average sound-evoked responses to the expected frequency were suppressed relative to the passive condition beginning with the first lever movement in a session and persisting throughout exposure to frequency deviants (Fig S2A,B), while neural responses to frequency deviants remained less suppressed throughout the experiment, which in some cases lasted up to 30 minutes and included hundreds of frequency deviants (Fig S2C). In a subset of mice, we presented a broad set of frequency deviants at half-octave intervals relative to the expected frequency and found that predictive suppression formed a notch that was specific to the frequency with which each mouse had been trained (Fig S2D). This notch was not present for mice prior to movement-sound coupling (Fig S2E), or at the end of one training session (Fig S2F,G), further highlighting that suppression of self-generated sounds in the auditory cortex is concentrated at a specific sound frequency learned over days of linked motor-sensory experience.

These experiments show that neural responses in the auditory cortex are suppressed for a self-generated sound that is irrelevant to a mouse’s behavior. But self-generated sounds are not always uninformative, and task relevance and reward association are known to enhance neural responses rather than suppress them^40,41^. We therefore asked whether predictive suppression was a core phenotype that always accompanied self-generated sounds, even those that are actively used to guide behavior. We trained a separate cohort of mice on a task variant where the sound played later in the movement (at the reward threshold), reasoning that this subtle change in the timing of the self-generated sound could make mice rely more strongly on the sound to guide their behavior (Fig 2A). Mice trained with the tone at the reward threshold experienced stronger empirical coupling between sound and reward than did mice trained to hear the tone early in their movement (Fig 2B). Consistent with this tighter coupling between sound and reward, these mice relied on the tone to guide their ongoing behavior, exhibiting extended reaches, prolonged lever trajectories, and delayed licking behavior when they experienced deviant sound frequency trials relative to interleaved control trials (Fig 2C,D). Mice reacted similarly when sounds were omitted (Fig S1G-L) and truncated their movements early when sounds were played early (Fig S1M-O), showing a bidirectional reliance on the sound to guide ongoing behavior. Despite this, neural recordings from mice trained on this alternative version of the task revealed that the modulation of auditory cortical responses to expected and unexpected self-generated tones was similar across the two training paradigms, with responses to expected sounds more strongly suppressed than responses to unexpected sounds (Fig 2E-H). These data indicate that the predictive suppression of expected self-generated sounds is a conserved phenotype that arises due to coupled motor-sensory experience regardless of behavioral relevance or reward association.

**Figure 2.**
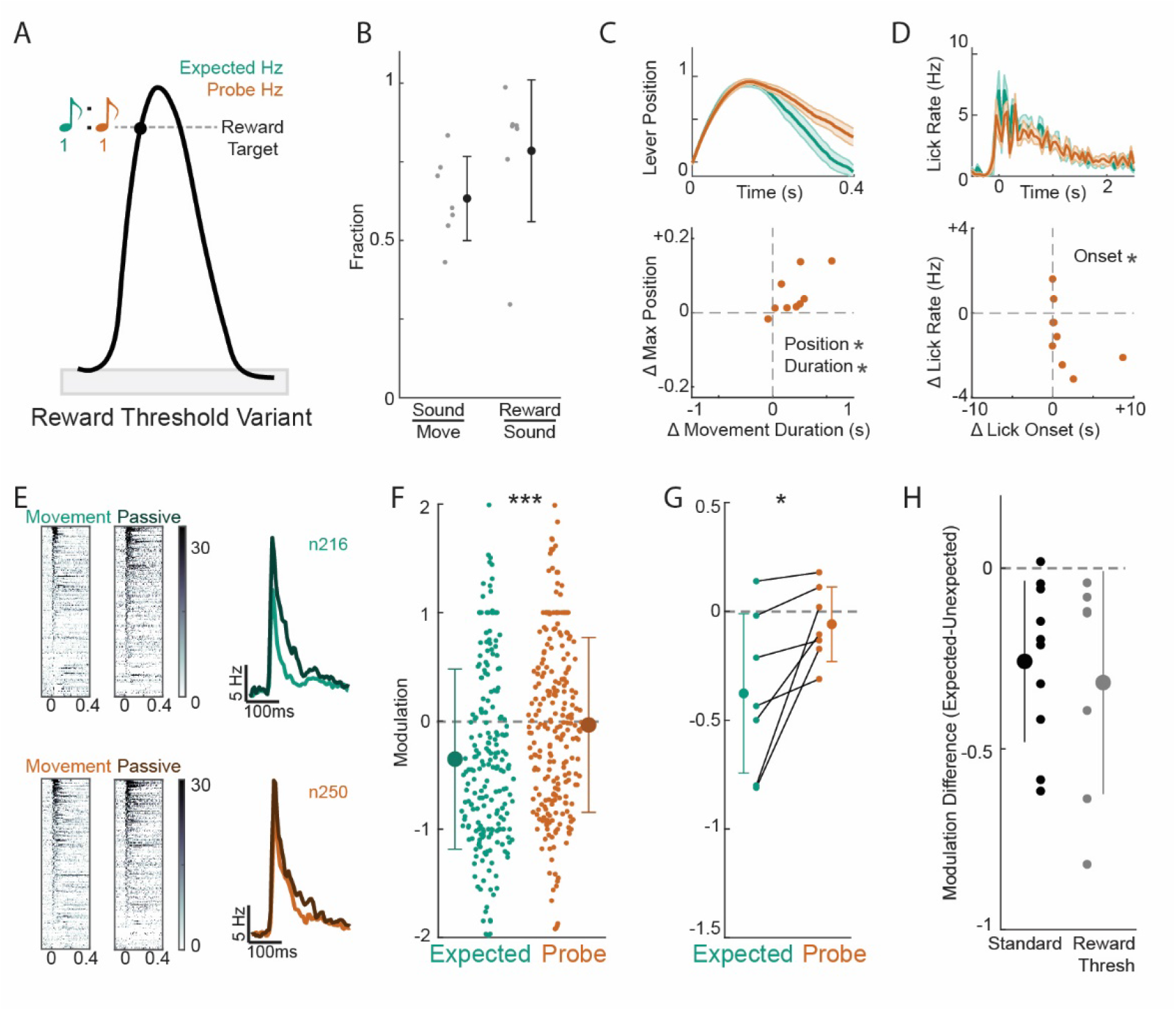
Frequency-specific suppression is present for self-generated sounds that are important to ongoing behavior. (A) Schematic of frequency-probe experiment where mice trained to expect the lever-generated sound at the reward threshold experience auditory feedback shifted by one octave on 50% of trials randomly interleaved. (B) Movement, sound, and reward coupling measured as the fraction of movements that generated sounds (crossed the sound threshold), and the fraction of sound-generating movements that generated rewards (obeyed inter-trial-interval and reached reward threshold)(N = 7 mice). (C) Global average lever trajectory +/− standard error across mice for interleaved expected-sound trials and frequency-deviant sound (top). Quantification (below) of change in movement duration (x-axis, seconds) and maximum lever position (y-axis, relative lever position) across interleaved expected and probe trials. (D) Global average pattern of mouse lick rate +/− standard error across mice for interleaved expected-sound trials and frequency-deviant sound (top). Quantification (below) of change in lick onset (x-axis, seconds) and lick rate (y-axis, Hz) for interleaved expected and probe trials. (E) Heat maps (left) of all neural responses (n=928, N=10) to the lever-associated sound (top, cyan) and frequency probe sound (bottom, orange) in the active condition (light, left) and passive condition (dark, right). PSTHs (right) of neural response to lever-associated sound (top, cyan) and frequency probe sound (bottom, orange) in the active condition (light) or passive condition (dark) averaged across all recorded neurons responsive to that sound in either the active or passive condition. (F) Modulation index of individual neurons comparing active and passive responses to the expected and probe frequencies. (G) Connected points display the average neural modulation of expected and probe frequencies for an individual animal. (H) Quantification of the average difference between the modulation of expected and probe frequencies for individual animals trained on the standard behavior (black, data from Fig. 1J) or the reward threshold behavior (gray). *p<0.05 ***p<.0005

Together, these findings reveal that mice trained to push a sound-generating lever develop a behavioral expectation for predictable self-generated sounds, and that expected sounds are subject to suppression in the auditory cortex that is stronger than sounds of an unexpected frequency. This frequency-specific suppression was learned, stable, and reflected movement-sound experience during training rather than tone relevance or reward association.

### Predictive suppression in auditory cortex is implemented at a specific time during a specific movement

These neural data reflect a commonly observed phenomenon – that expected sounds experience suppression relative to unexpected sounds in the auditory cortex – and also show that this modulation is stable over time and is linked to motor-sensory experience during training rather than tone relevance or reward association. These properties align with the implementation of a movement-based prediction in auditory cortex ^3,14^, but a critical outstanding question is whether this frequency-specific modulation is organized temporally around a specific movement. The separable and prolonged movements in our lever paradigm enabled us to measure the time-course of neural modulation within and around individual movements.

We first asked whether frequency-specific suppression is restricted to periods of lever movement or represents a slower modulation related to the mouse’s general behavioral state. In a separate group of mice trained on our standard lever task, we recorded neural responses to lever-generated sounds while playing randomly timed lever-associated and probe frequency tones in the background, uncoupled from behavior (Fig 3A). By chance, many of these background tones occurred just prior to a lever movement, providing an opportunity to explore the time course over which frequency-specific suppression is applied before and during behavior. Neural responses to background tones heard prior to the onset of lever movements displayed minimal modulation compared to passively heard tones for both the lever-associated and probe frequencies (Fig 3B). Predictive suppression of the expected frequency emerged during the lever press, with neural responses to the expected frequency more strongly suppressed than neural responses to the unexpected frequency (Fig. 3B). These data show that both general suppression and frequency-specific suppression are implemented only within the temporal confines of individual lever movements, despite the fact that lever movements often occur within seconds of one another.

**Figure 3.**
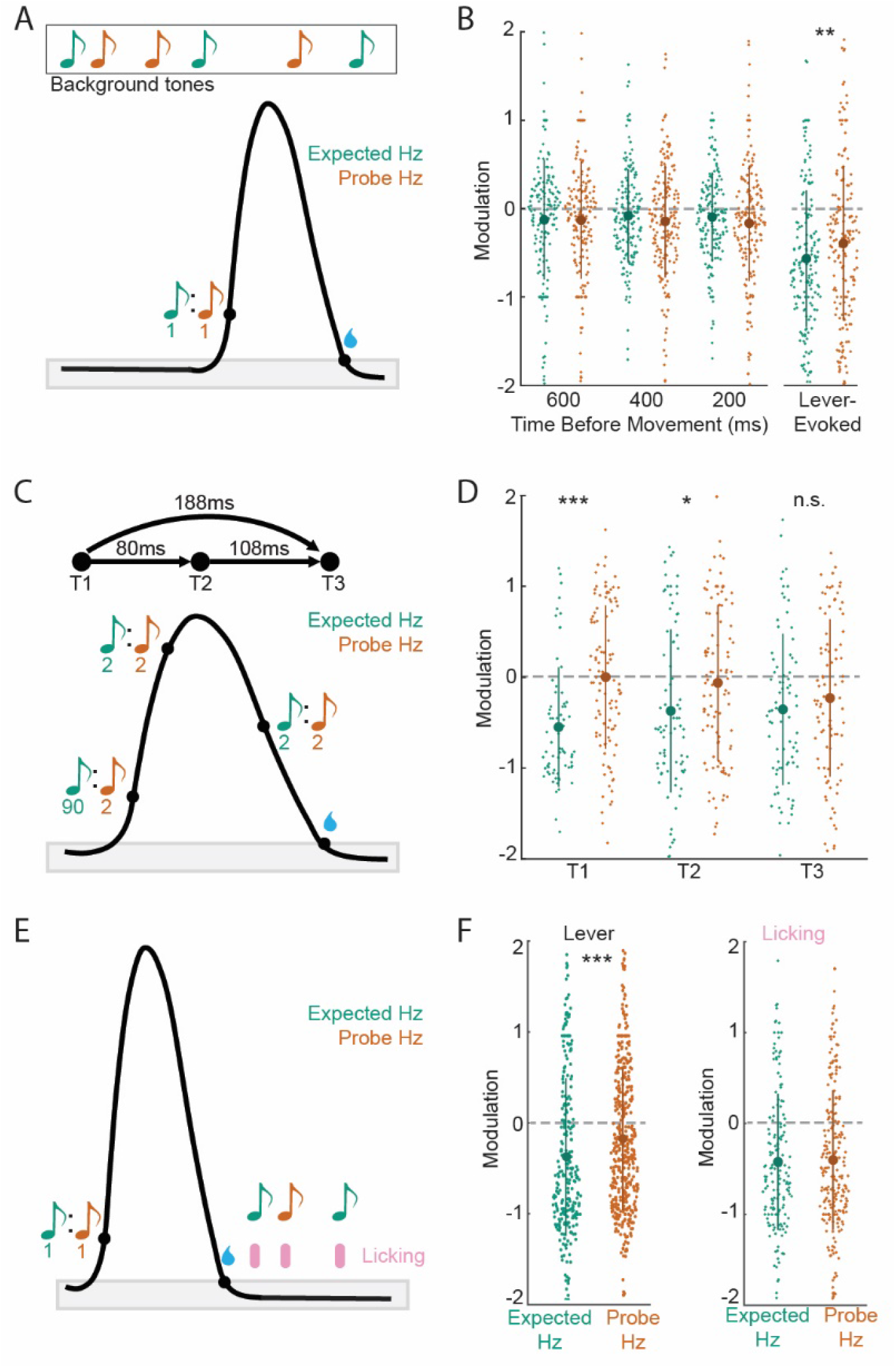
Frequency-specific suppression is confined to a specific time point within the sound-associated movement. (A) Schematic of timing probe experiment, with mice experiencing randomly played background tones in addition to standard and frequency-probe lever-generated sounds early in movement (N=4). (B) Average modulation of neural responses to lever-associated (cyan) and probe (orange) tones heard during lever movement, or during 200ms bins prior to lever movement onset, starting at the time listed on the x-axis. Modulation values are calculated relative to passively heard tones at experiment conclusion. (C) Schematic of within-movement probe experiment (bottom) where expected and unexpected frequency tones are played at the expected position, at a point near the apex of movement, or in the middle of the return phase (N=4). 90% of sounds were the expected frequency at the expected position, with the remaining 10% distributed across the other 5 frequency/position combinations. Timing (above) shows the difference between time points, averaged across animals. (D) Average modulation of neural response to the expected (cyan) and probe frequency (orange) tones at three different time points. Neurons are included at each time point if they have a significant response to the corresponding tone at that time point or in the passive condition. (E) Schematic of movement-specificity experiment where mouse licking triggers closed-loop generation of the lever-associated frequency or probe frequency, randomly interleaved (N=6). (F) Average modulation of the expected and frequency probe sounds in this subset of animals when heard at the expected lever position (left) or when triggered by licking (right). *p<0.05 **p < 0.005 ***p<.0005

Having established that frequency-specific suppression was not present prior to movement onset, we next asked whether this suppression displayed temporal specificity within the time course of each lever press. In a separate group of mice trained on our standard lever task, we recorded neural responses to self-generated sounds played at unexpected positions on a small fraction of trials. This allowed us to measure the strength of frequency-specific suppression at three separate points within the movement for each animal: the expected position, late in the outbound phase, and the middle of the return phase (Fig 3C). Consistent with our standard frequency-probe experiments, we observed strong frequency-specific suppression when sounds occurred at the expected position early in movement (Fig 3D). The strength of this frequency-specific suppression diminished as sounds were played farther from the expected position. Sounds played late in the outbound phase, delayed on average by 80ms, displayed weaker frequency-specific suppression, and sounds played during lever return displayed no significant difference between the modulation of the two sounds (Fig 3D, 188ms average delay). These data show that frequency-specific suppression has temporal specificity on the scale of 10s to 100s of milliseconds, revealing a previously unknown level of precision that matches the ethologically relevant time scale for error detection ^38,42,43^.

Finally, we tested whether this expectation-based modulation is specific to the movement that was paired to sound during training, or whether it generalized to other movements, especially those that are temporally proximal to the sound-associated movement. We trained mice on our standard sound-generating lever task and then performed neural recordings while delivering tones triggered by licking movements in addition to the lever (Fig 3E). This paradigm allowed us to present tones, including the lever-associated frequency, during two unique movements that occurred within hundreds of milliseconds of one another. In contrast to the frequency-specific, predictive suppression observed during lever movements, there was no frequency-specific suppression to tone-evoked responses during licking (Fig 3F). The lack of predictive suppression for tones heard during licking highlights the exclusive relationship of predictive suppression to the trained movement even over short time scales and confirms the temporal-specificity of the lever-based prediction.

Together, these data establish that the frequency-specific modulation of expected, self-generated sounds results from a movement-based prediction implemented over short time scales at a specific time-point during a specific movement.

### Movement signals in auditory cortex reflect expectations for the frequency and timing of self-generated sounds

Our results show that auditory cortical responses to expected, self-generated sounds experience a pattern of predictive modulation that is learned, stable, and implemented with sub-movement specificity around an individual physical action. There are multiple lines of evidence suggesting that the implementation of this prediction might happen locally. Frequency-specific suppression of self-generated sounds is absent in the sensory thalamus ^28,29^, the auditory cortex receives long-range input from motor-related structures that are known to be involved in modulating auditory cortical activity during behavior ^4,28^, and representations of orofacial movements have been found in some cortical layers ^44^. Despite the known motor-related modulation of auditory cortex, it remains unknown how prevalent movement signals are in the auditory cortex, and whether they are specific enough to encode an expectation for the timing and acoustics of a self-generated sound.

To investigate the properties of movement signals in the auditory cortex, a group of animals trained on the standard behavioral task also experienced a small fraction of trials where sound was omitted in addition to timing and frequency deviant trials (Fig 4A, see methods). These trials provided an opportunity to analyze activity in the auditory cortex that was related strictly to movement and expectation, and we began by identifying all neurons that had significant responses in a 50ms window surrounding different time points throughout the movement (See Methods). We found a large population of neurons that were active during movement in the absence of auditory feedback (36%, Fig S4A) and plotted their average activity aligned to the time of expected sound onset (Fig 4B). Average population activity began to ramp up roughly 200ms prior to movement onset and peaked almost exactly at the time of the expected tone, with activity then decaying for roughly 400ms, which corresponded roughly to the time of the movement duration (Fig S4B,C). To ensure that the peak we observed at the time of expected auditory feedback was not an artifact of aligning neural activity to the time of the expected sound, we aligned the activity of the same neural pool to different positions within the movement and found that peak average firing always corresponded to the time of the expected sound (Fig 4D).

**Figure 4.**
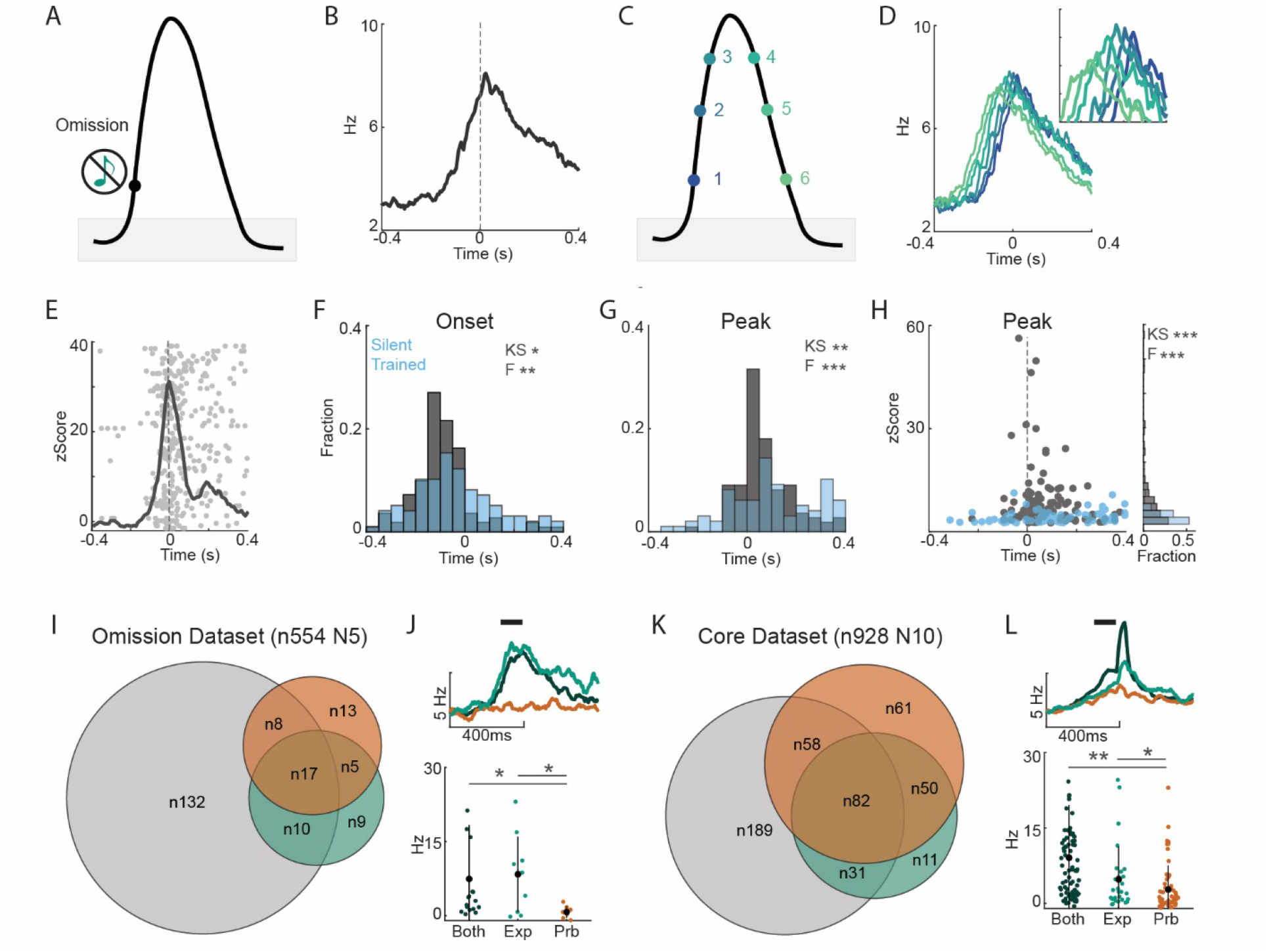
Movement signals in the auditory cortex display an expectation for the timing and frequency of self-generated sounds. (A) Schematic of omission trials occurring in a subset of mice trained on the standard sound-generating lever task (N5). (B) Average population PSTH of all neurons responsive to any of 6 positions throughout movement (see methods), aligned to the time of expected sound. (C) Schematic showing the 6 positions throughout movement, corresponding to the time points used to identify movement-responsive neurons. (D) PSTHs of entire movement-responsive pool aligned to each of the time points shown with corresponding color in (C). (E) Average z-scored response of an individual movement-responsive neuron over a raster plot of action potential time across trials, aligned to the time of lever-initiated sound onset. (F) Probability histogram of movement-signal onset time on omission trials for individual neurons in standard trained mice (gray) and mice trained on the same task but in the absence of sound (blue). Distributions were tested for similarity using the KS test and the f-test. (G) Probability histogram of movement-signal peak time on omission trials for individual neurons in standard trained mice (gray) and mice trained on the same task but in the absence of sound (blue). (H) Intersection of peak time and strength of neuron movement signal (z-score) for standard (gray) and silent-trained (blue) mice, with histogram of movement signal strength (right). (I) Venn diagram showing the intersections of movement responses and sound responses to the expected and probe frequency tone across the omission dataset population. (J) PSTH (top) showing the movement activity on omission trials of neurons responsive to the expected sound (cyan), the probe sound (orange), or both sounds (dark cyan). X-axis notch indicates the time of expected sound. Quantification (bottom) of neural response rate in the 100ms prior to the expected tone time for each population. (K,L) Same as (I,J), but for neurons in the standard dataset that lacked omission trials. X-axis notch (top) indicates the time of sound. *p<0.05 **p < 0.005

Having identified neurons active during lever movement in the absence of sound in the auditory cortex, we examined movement-related activity across individual neurons. For each neuron, we determined the onset time and peak time of movement-related activity, as well as the magnitude of movement-related signal strength (z-score) (Fig 4E). Like the average population response, many individual neurons began responding about 300-400ms prior to the time of sound, with other neurons initiating their responses throughout the course of the movement (Fig 4F). The peaks of movement-related activity for individual neurons were not homogenous but instead were concentrated around the time of the expected sound (Fig 4G), and neurons with activity peaks around the time of the expected tone also had the largest movement-related changes in firing rate (Fig 4H, Fig S4D). This tight alignment of movement signal timing and strength with the time of expected sound makes a strong case that these neurons function as an expectation signal in the auditory cortex. To confirm that this pattern constitutes a learned expectation for the timing of the expected tone, we performed an identical analysis in a cohort of mice trained to push a lever in the absence of sound (See FigS2E). These mice made lever movements of similar magnitude and duration and had a similar fraction of auditory cortex neurons active during their reaches (Fig S4A,B). In contrast to mice trained with a sound-generating lever, the peaks of movement signals in mice trained on a silent lever were distributed more homogenously throughout the lever movement, while the magnitudes of movement-related signals were weaker on average without a clear temporal organization (Fig 4F-H. These findings indicate that experience with a sound-generating movement shapes movement-related signals in the auditory cortex to become concentrated at the time of an expected self-generated sound.

We next asked whether movement-related signals in the auditory cortex also reflected an expectation for the specific acoustic features of a self-generated sound. We segregated the movement-responsive neurons described above into groups based on their frequency tuning. Although we found roughly equal numbers of neurons with movement-related activity that were tuned to the expected frequency and probe frequency (Fig 4I), the magnitude of movement-related activity was significantly stronger for neurons that were tuned to the expected frequency (Fig 4J). To confirm that these results were consistent with our larger data set that focused on frequency deviants and lacked omission trials, we identified neurons with movement-related signals specifically in the time just prior to the self-generated tone (Fig S4A-C). As in our omission data set, we found that a similar fraction of neurons tuned to the expected or unexpected sound contained movement responses (Fig 4K), and that neurons tuned to the expected sound experienced the strongest movement signals (Fig 4L).

Taken together, these data show that movement-related activity in the auditory cortex – observable when acoustic feedback is omitted – reflects a learned expectation for the timing and frequency of a self-generated sounds. These activity patterns, present in average population activity as well as many individual neurons, signal expectation at a level of specificity commensurate with the temporally precise modulation of self-generated sounds observed during lever movements.

### Distinct laminar patterns of expectation signals, sound representations, and violation signals

Average neural activity patterns in the auditory cortex reveal signals related to movement, expectation, and sound during the generation and processing of self-generated sounds, but with substantial heterogeneity across the recorded population. In the cerebral cortex, depth from the pial surface is a principal organizing feature, with neurons in different layers possessing distinct intrinsic properties, gene expression, and local and long-range projection patterns. To determine whether predictive signals displayed organization within the auditory cortex, we investigated the distribution of movement, expectation, and sound signals across cortical layers.

First, we measured the properties of movement-related neurons throughout the cortical column by assigning each neuron a putative cortical depth relative to consistently observed current source density features (Fig 5A, See Methods). Neurons with movement-related activity were distributed across cortical layers but were most abundant in L6, which contained the largest number and fraction of movement-responsive neurons (Fig 5B). Plotting the onset time and peak time of each neuron’s activity by cortical depth showed that the earliest movement signals were found deep in the cortex (Fig 5C), where the onset and peak of movement signals tiled a broad range of times over the course of movement (Fig 5C,D). In contrast, more superficial neurons typically became active shortly before the time of expected sound and peaked shortly thereafter (Fig 5C,D). Measuring the intersection of movement signal timing and response strength revealed that L6 contained many weak movement-related signals positioned throughout the duration of the movement in addition to strong signals centered on the time of expected sound (Fig 5E). Neurons outside of L6 had movement signals with a similar range of amplitudes, but which were more closely aligned to the time of expected sound (Fig 5E, f-test, L6 vs non_L6 p = 0.003). The preponderance of signals throughout the time-course of movement in deep auditory cortex is consistent with L6 neurons encoding orofacial behaviors and with anatomical connections from motor-related regions onto L6 excitatory neurons ^2,44^. The abundance of strong movement signals aligned to the time of expected sound, which dominate movement responses in more superficial layers, suggest that aspects of movement related to expected auditory outcomes are transmitted broadly across cortical lamina.

**Figure 5.**
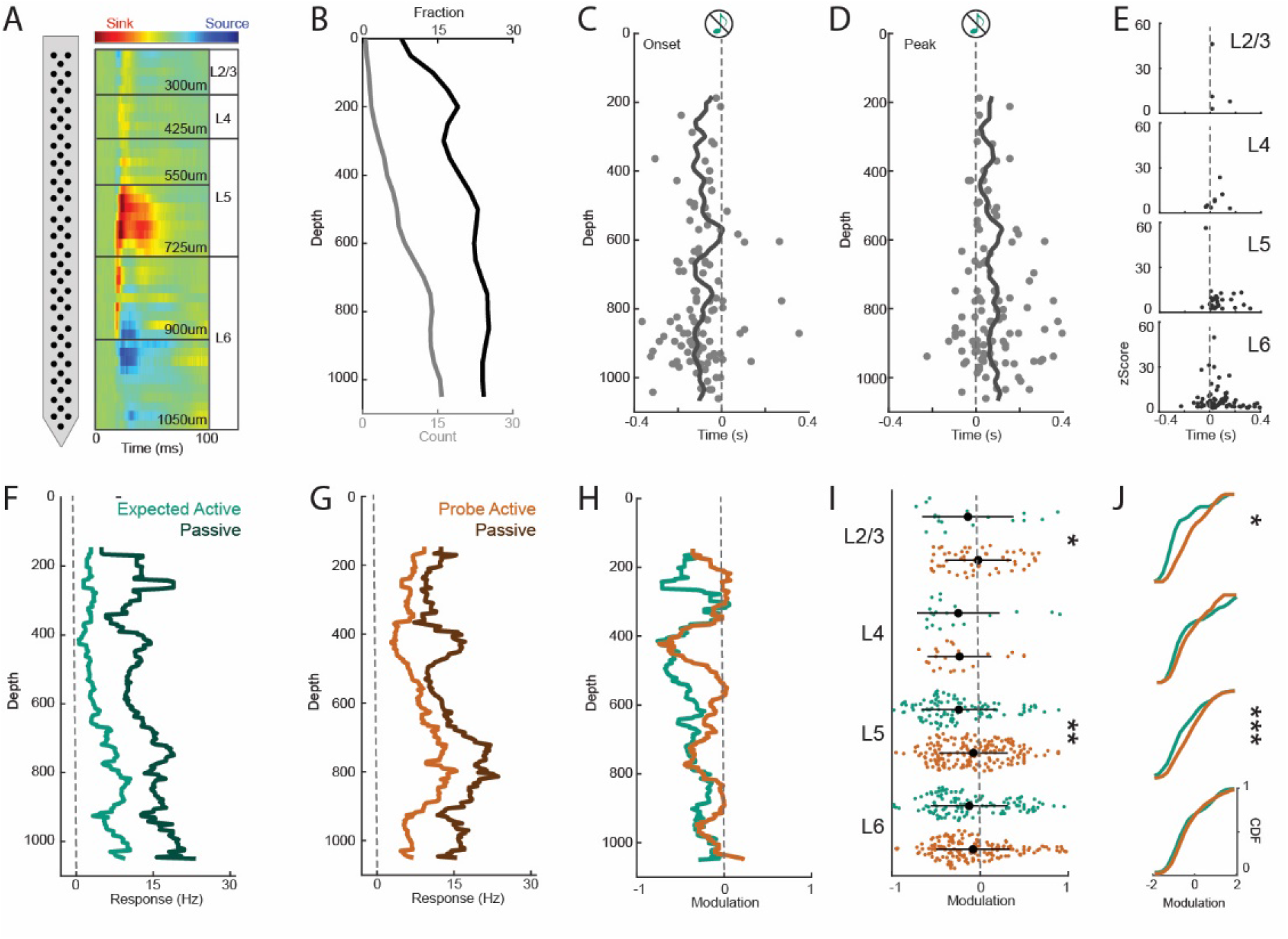
Laminar pattern of movement, sound, and expectation signals. (A) Schematic showing the calculation of relative cortical depth and laminar assignment based on current-source-density (CSD) features (see Methods for details). (B) Moving average histogram showing the number (gray) and fraction (black) of neurons responsive during lever movement when sound is omitted (see Fig 4A,B). (C) Onset time of movement responsive neurons plotted against cortical depth with moving average line (averaged across 50um). (D) Same as (C) but for movement response peak. (E) Intersection of peak time and z-score strength of neuron movement signals broken down by cortical layer. (F,G) Moving average (across 50um) of individual neuron responses to the expected frequency sound (F, cyan) or probe frequency sound (G, orange) heard in the active (light) or passive (dark) context plotted by neural depth. (H) Moving average of neural modulation to expected and probe sounds across cortical depth (overlapping 50um bins). (I,J) Radial modulation comparing neural responses in the active and passive condition for expected (cyan) or probe (orange) sounds in individual layers (I) and corresponding cumulative distribution histograms of neural modulation for each layer (J). * p < 0.05**p < 0.005 ***p<0.0005

We next examined the laminar distribution of sound-evoked responses by comparing the magnitude of suppression of the expected versus the probe frequency at different cortical depths. Frequency-specific suppression was strongly concentrated in L5 and, despite having a smaller sample size, in L2/3 (Fig 5F-J). In both L2/3 and L5, the magnitude of neural responses to probe frequencies were minimally suppressed, while neural responses to the expected frequency were strongly suppressed. In contrast to L2/3 and L5, the average level of movement-based modulation was comparable for both expected and unexpected tones in L4 and L6 (Fig 5F-J). While sound responses in L4 neurons were strongly suppressed, neurons in L6 on average experienced minimal suppression and had the largest average sound-evoked firing rates in the cortex (Fig 5F,G; Fig S5).

The laminar distributions of movement signals and responses to expected and unexpected sounds reflect a division of labor where different aspects of predictive computation are concentrated in distinct layers.

### Predictive processing signals are carried by distinct functional groups in auditory cortex

Finally, we asked how signals related to predictive processing were distributed across individual neurons in the auditory cortex. We started with a manual analysis focusing on neurons that, in spite of population-level suppression during movement, retained a significant response to at least one of the self-generated sounds (expected frequency or probe frequency). A large population of neurons, concentrated in L6, responded ubiquitously to both the expected and probe sounds whether they were self-generated or heard passively (8%, Fig 6A,E,F). A smaller group of neurons, also strongly represented in L6, responded to both sound frequencies but only during movement (4%, Fig 6B,E,F), displaying a movement-enhanced phenotype that was in contrast to the average phenomenon of suppression observed during movement (see Fig. 1I) and consistent with the presence of movement-related signals in deep layers of auditory cortex (see Fig. 5B-E).

**Figure 6.**
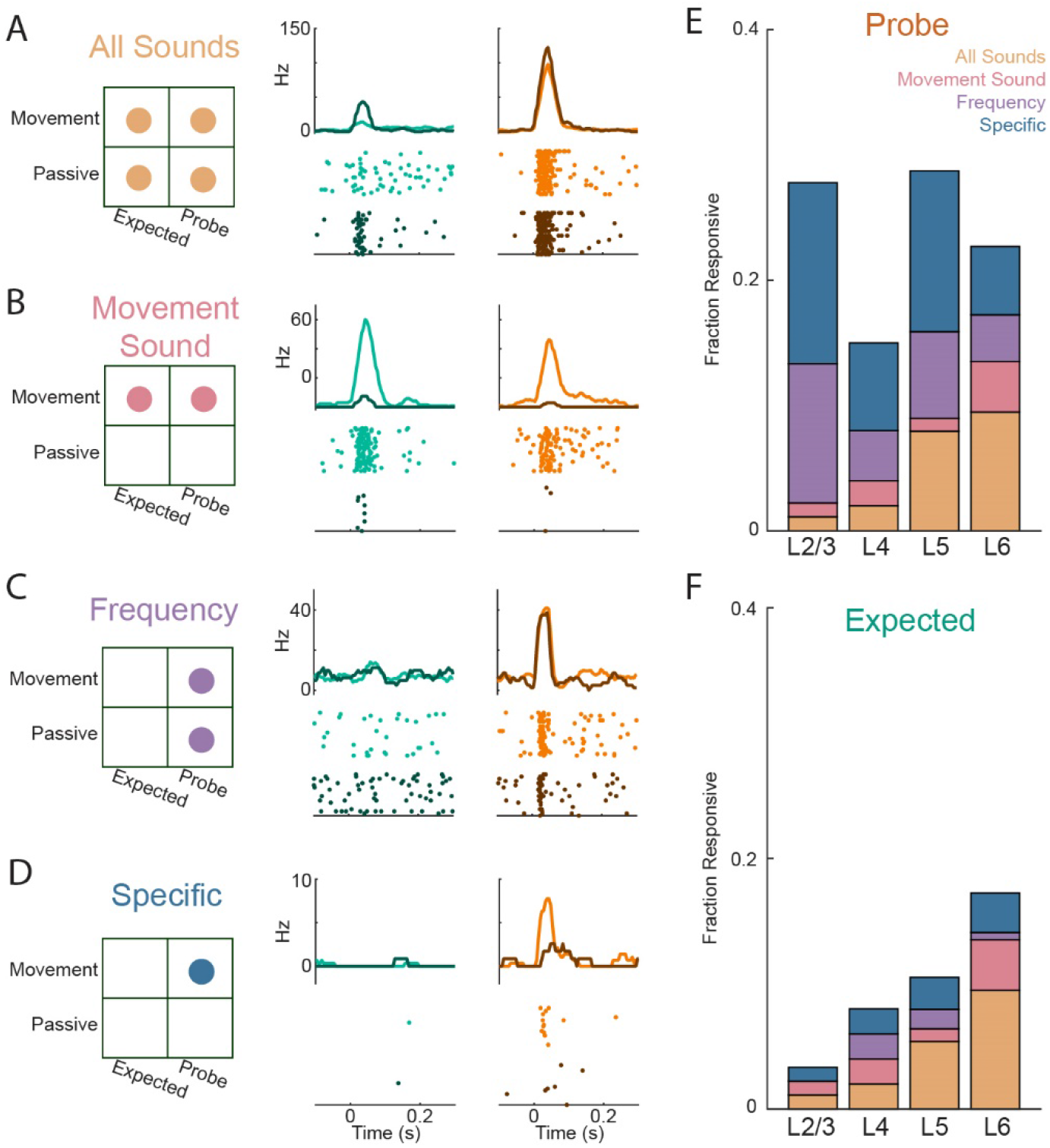
Divergent processing of expected and unexpected self-generated sounds across individual auditory cortex neurons. (A-D) Schematic and neural activity of a sample neuron exemplifying cells responsive to all sounds across all conditions (A), responsive to both sounds only during movement (B), responsive to a specific frequency in both the active and passive context (C), and responsive specifically to either the expected or probe frequency during movement (D). Circles in schematic (left) indicate responsiveness across trials (p < 0.05) to a given combination of tone identity (expected, probe) and context (movement, passive). Raster plots (bottom right) show action potentials across trials for expected (cyan) and probe (orange) frequencies heard in the active (light) or passive (dark) condition, and PSTH (top right) show average neural response of the neuron across trials. (E-F) Fraction of neurons in each layer responsive to the self-generated probe sound (E) or self-generated expected sound (F), with stacked bars representing the fraction of neurons that fit into each of the functional groups in A-D.

We next focused on neurons that were responsive to only one of the two self-generated sounds. Many neurons, especially in L2/3 and L5, responded to the unexpected tone when it was heard passively and when it was self-generated (9%, Fig 6C,E). In contrast, far fewer neurons responded to the expected tone in both behavioral conditions (2%, Fig 6F). This near absence of neurons that respond to the expected tone in the passive condition and retain a response when it is self-generated mirrors the larger core phenotype of strong movement-related suppression applied to a single, expected frequency. Finally, we identified a large population of neurons that responded only to the unexpected frequency tone and only when it was heard during movement (10%, Fig 6D,E). These neurons were particularly abundant in L2/3 and L5 and signal a specific mismatch between expectation and experience. In contrast, very few neurons responded only to the expected tone heard during movement (3%, Fig 6F), revealing a striking asymmetry in the way individual neurons respond to expected and unexpected self-generated sounds.

Our analysis of individual neuron and population-level activity has been largely hypothesis-driven and has uncovered a variety of neural signals related to the implementation and outcome of movement-based predictions in auditory cortex. These include strong frequency-specific suppression of sound responses in L2/3 and L5, general movement-evoked suppression particularly in L4 and L6, neurons in L2/3 and L5 that are active only following a violation of expectation, and both general and specific movement signals. We conclude our study by performing machine-learning based clustering of neurons based on their movement- and sound-related activity to determine whether predictive processing signals are identified with an unbiased analysis method and to gain insight into their distribution across individual neurons. We characterized each neuron by creating a vector of activity variables that encompassed baseline, sound-evoked, and movement-evoked activity (Fig. 7A, see Methods). We applied a k-medoids clustering algorithm^45^ to separate neurons into discrete groups based on similar response features and we visualized the clustering results by plotting each neuron in a low-dimensional projection of our feature space, color coded by cluster identity (Fig 7B).

**Figure 7.**
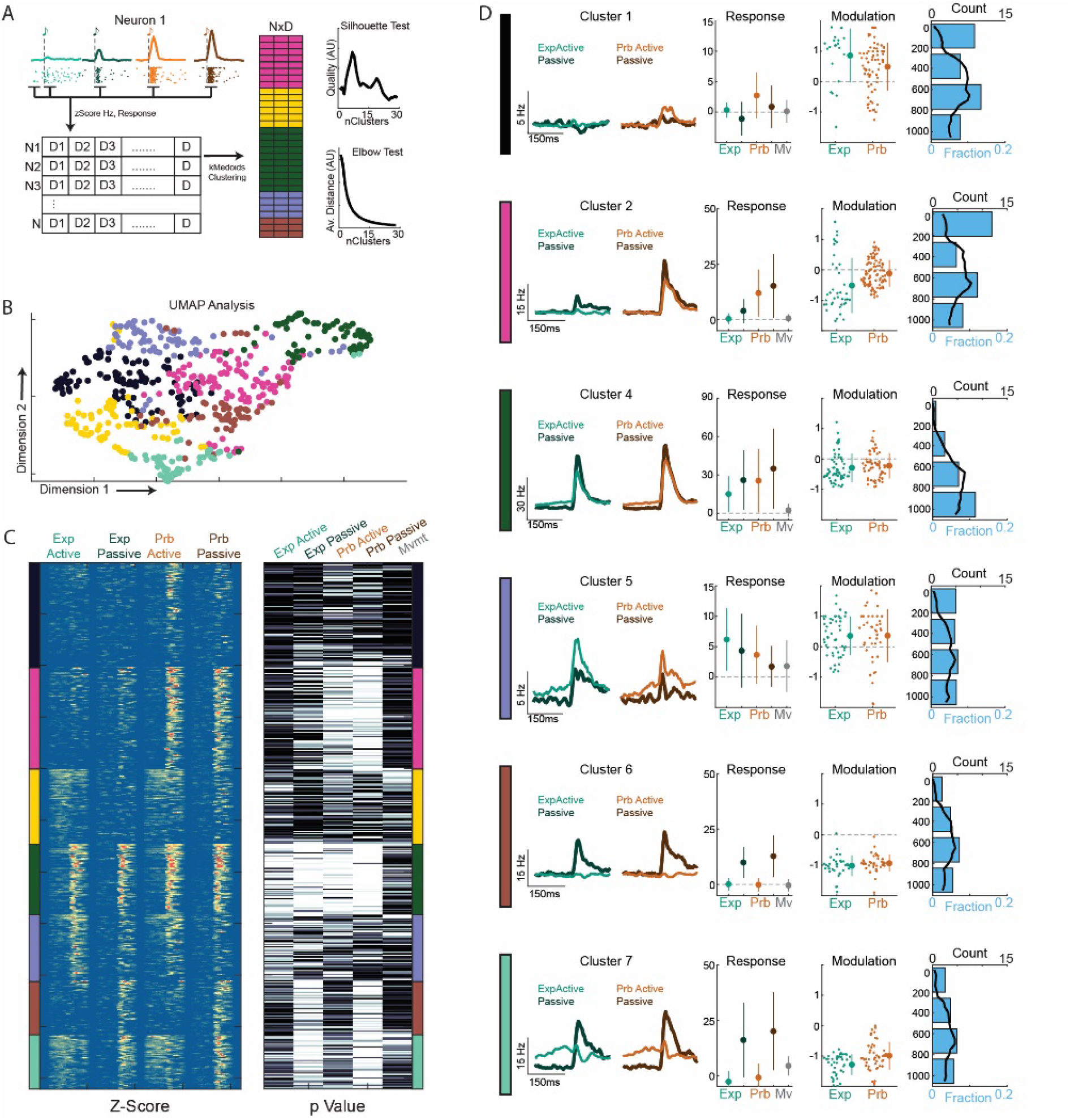
Movement, expectation, sound, and error signals are distributed across distinct functional groups. (A) Schematic (left) of clustering process, where individual neurons are characterized by their responses to different task variables and then grouped using k-medoids clustering based on these dimensions (Core dataset, N10, see methods). Silhouette test (top right) showing the quality of clustering for different cluster numbers (k-values), and elbow test (bottom right) showing the average distance of all data points to group center as a function of cluster number. (B) Visualization of cluster groups (k = 7) by plotting each neuron in a low-dimension projection of our feature-space generated by UMAP and color coded by cluster identity. (C) Neuron response patterns sorted by cluster group (designated by color), shown as the z-scored activity pattern over time to each sound type, and as response p-values following sound onset (0-60ms post sound vs 100ms pre-sound) or prior to sound onset for movement signals (100ms prior to sound vs 400-500ms prior to sound). (D) Functional characterization of select cluster groups. From left to right: PSTHs of average neural response to expected (cyan) and probe frequency (orange) sounds in the active (light) and passive (dark) conditions. Note different Y axis scales. Quantification of average response strength across all neurons to each sound type and movement signals as measured in (C). Quantification of modulation of each neuron to expected and probe frequency sounds, with neurons included if they have a significant response to the given tone in either the active or passive condition. Right, quantification of neuron counts by depth (top, black) and fraction of all neurons in each layer (bottom, blue).

The clustering output, visualized as sorted neural response heatmaps and aggregate response patterns for each group, identified populations of functionally similar neurons that accounted for many of the signals observed during our manual analysis (Fig 7C,D). First, we identified two populations of neurons concentrated in L2/3 and L5 that reflect the predictive processing of self-generated sounds without displaying any movement-related signals. One group was comprised of generally sound-responsive neurons that experienced preferential suppression of expected, self-generated sounds (Cluster 2). The other group consisted of prediction-error neurons, which responded only to self-generated sounds that violated frequency expectations (Cluster 1). A third group of sound-responsive but not movement-responsive neurons, dispersed across cortical depths, displayed complete movement-based suppression of sound responses regardless of frequency (Cluster 6). Movement signals leading up to the self-generated sound were distributed across 4 clusters, including a group of neurons concentrated in L6 that displayed strong responses to both sound frequencies in both the self-generated and passive conditions (Cluster 4), a group that displayed enhanced responses to self-generated sounds regardless of frequency (Cluster 5), and a group that experienced abrupt suppression following the onset of self-generated sounds (Cluster 7).

Because our primary dataset doesn’t allow us to observe movement signals in the absence of sound, we performed an identical clustering analysis on our dataset that included sound omissions in addition to frequency deviations (Fig S7A). Despite being a smaller dataset and employing a different probing strategy, clustering yielded many functional groups that were virtually identical to those observed in our primary data set (Fig S7C,D), including predictive suppression neurons and prediction error neurons in L2/3 and L5 (Cluster S6,7,8). This dataset allowed us to include the activity of neurons at the time of the expected sound on omission trials, which provided added resolution of movement-responsive populations. As above, a group of neurons in deep layers displayed both pre-sound movement signals and sound responses (Cluster S6), and the inclusion of omission trials shows that these neurons lack a specific peak at sound time in the absence of sound. Two groups of movement-responsive neurons were not responsive to passive sounds, ramped up their firing during lever movements, and exhibited a peak at the time of expected sound regardless of the presence or absence of auditory feedback (Cluster S5,9). These populations were distributed across cortical depths and distinguished by the strength of their movement-related activity, with the smaller group (3% of neural population) displaying particularly strong and temporally precise activity at the time of expected sound (Cluster S9).

These unbiased clustering analyses recapitulate the core findings from our manual analysis: that expected and unexpected sounds are processed differently in L2/3 and L5 due to separate groups of predictive suppression and prediction error neurons; that there are abundant movement signals in deep layers of the auditory cortex; and that a smaller population of neurons distributed across cortical depths has strong movement-related responses centered precisely at the time of expected sound. Together, our experiments reveal a consistent and structured functional organization of movement, expectation, sound, and error signals during the implementation of precise movement-based predictions in the auditory cortex.

## Discussion

Our results show that frequency-specific suppression in the auditory cortex arises from a stable, learned, and specific movement-based prediction that is implemented over short time scales with within-movement temporal specificity. This suppression is applied to expected sounds regardless of their behavioral relevance, is concentrated in L2/3 and L5 and is preceded by the onset of learned movement signals that encode an expectation for the frequency and timing of an expected sound. Examining the distribution of movement and sound signals across individual neurons revealed predictive suppression and prediction error neurons in L2/3 and L5, general movement-related signals in L6, and expectation neurons distributed across cortical layers that precisely mark the time of expected sounds. The confluence of movement, sound, expectation, and error signals in the auditory cortex reveals the implementation of precise movement-based predictions that may enable animals to identify deviant acoustic feedback and, when appropriate, adjust subsequent behavior^5^.

Multiple lines of evidence in our study indicate that frequency-specific suppression in the auditory cortex is driven by a learned motor-sensory prediction rather than short-term adaptation, arousal, or general movement-based suppression^2,22,37,46,47^. Because the expected and unexpected frequency tones were interleaved with equal likelihood during electrophysiology, our experimental design precluded frequency-specific suppression caused by short time-scale effects like stimulus-specific adaptation^37,46^. Instead, suppression was only visible after multiple days of training, and was present immediately at behavior onset and stable over tens of minutes in trained mice. Importantly, we observed comparable levels of frequency-specific suppression even during a task variant where the tone is highly predictive of reward and mice actively rely on the tone to guide their ongoing behavior. Together, these findings show that frequency-specific suppression of self-generated sounds is a stable, learned, and intrinsic outcome of motor-sensory predictions.

Although the phenomenon of frequency-specific suppression has been studied using augmented reality paradigms in the past, previous experimental designs have not been able to establish that this modulation is directly linked to individual movements or to identify expectation signals linking movement and sound. Prior behavioral paradigms have lacked a continuous readout of a prolonged and isolated movement (e.g. licking or button pressing) while others have coupled self-generated sounds onto a more abstract readout of a continuous behavior such as locomotion velocity^6,28,29^. Our experimental paradigm allowed us to present a self-generated sound at an arbitrary time within a continuous behavior, to finely adjust the timing of self-generated sounds within the movement time-course, and to monitor in real time two distinct but temporally proximal behaviors. While not technologically groundbreaking, this approach was transformative in allowing us to test the specificity of movement-based predictions along multiple dimensions (stimulus, timing, and movement) in a single behavioral paradigm. We found that the suppression of expected, self-generated sounds was only engaged during sound-producing movements, was specific to the movement coupled to sound during training, and had within-movement temporal specificity on the order of tens to hundreds of milliseconds. These predictions possess a degree of movement and temporal specificity consistent with the requirements of ethologically relevant error detection and matches the dysfluency that accompanies delayed auditory feedback in humans and songbirds^42,43^.

Our experimental paradigm also allowed us to examine the time-course of non-sound information in the auditory cortex. Movement-related modulation of sensory cortex has been widely reported, and some types of modulation are influenced by direct transmission of motor information to the auditory cortex ^4,15,17,19,21,22^. Recently, motor signals themselves have been observed in auditory cortex neurons during orofacial movements^44^. However, little was known about the prevalence of movement signals in the auditory cortex, and whether movement signals are able to encode predictive information about the sensory outcomes of movement. By recording neural activity on interleaved sound omission trials, we were able to observe movement-specific movement-related expectation signals in auditory cortex neurons. Roughly one third of auditory cortex neurons were responsive at some point during lever movement, both in mice trained with acoustic feedback and mice trained to perform the same task but in silence. However, mice trained to expect a self-generated sound had movement responses that were larger and peaked at the time of the expected sound, with the strongest responding neurons having peaks most closely aligned to the time of the expected sound. Meanwhile, movement signals in mice trained to push a quiet lever were weak and distributed homogenously throughout movement. In addition to this temporal expectation, neurons tuned to the frequency of the expected self-generated sound had stronger movement-responses than neurons tuned to other frequencies. Taken together, these movement-related signals reveal the presence of learned expectations in the auditory cortex that were specific for both timing and stimulus identity. The exact origin of these movement signals is not known, but they could arrive via one or more of the motor cortex, the basal ganglia, or the higher-order thalamus, which processes both movement and sensory information^2,4,24,44,48–51^. Regardless of origin, the presence of movement-related signals that are specific to the time and frequency of a self-generated sound may reflect an experience-dependent transformation of movement signals from a motor-centric coordinate frame into sound-centric coordinates^24,52^. It remains to be established whether motor signals arrive at the auditory cortex in sound-centric coordinates or whether they are transformed locally within the auditory cortex.

Finally, our recording paradigm allowed us to establish that movement, expectation, and sound signals were not uniformly distributed across cortical layers or individual neurons. Weak movement responses were most abundant in L6 and were found in both sound-responsive and non-sound-responsive populations^44^. Strong movement-signals that peaked precisely at the time of expected tone emerged from our unbiased clustering as a distinct functional group and were distributed across cortical depths rather than being concentrated in L6. Conversely, frequency-specific suppression was strongly concentrated in L2/3 and L5, but not in L6 and L4. Many neurons in L4 and L6 receive strong excitatory input from the primary sensory thalamus which does not display frequency-specific suppression^28,29,50,53–58^, consistent with predictive modulation arising *de novo* at the level of the cortex. The difference in sound responses in L2/3 and L5 were driven by two functional groups of neurons: one that exhibited within-neuron frequency-specific suppression of sound responses, and a group of neurons that responded specifically to unexpected sounds but were otherwise not driven by sound. These prediction error neurons were abundant in L2/3 and L5, making up 14% of all neurons. Visual flow mismatch neurons, a conceptually similar functional group, have been observed in L2/3 but not L5 of primary visual cortex^3,14,33–35^. These prediction error signals provide a potential neural substrate for behavioral reactions to deviant acoustic consequences and may play a role in auditory-guided motor learning in more complicated behaviors^11,59,60^. Despite the average suppression of expected sounds, 15% of the population still responded to the expected, self-generated tone in some capacity. Many of these neurons were ubiquitously sound-responsive cells in L6 that responded to both sounds in both behavioral conditions. Given that some responses to the expected sound are retained and that mice experience suppression even when they actively rely on a tone to guide their behavior, it is likely that predictive suppression alters but does not abolish the representation of expected sounds in the auditory cortex. Biologically, smaller pools of responsive neurons may represent expected sounds more efficiently, and refined representations may increase the amount of information that can be represented by a finite neural pool^61–65^.

These findings reveal distinct functional groups of auditory cortical neurons with activity patterns reflecting movement, expectation, sound, and prediction error. This paves the way for future experiments, aimed at understanding predictive modulation at a mechanistic level, that map these functional groups onto biologically defined cell types. Our results also set the stage for studying how synaptic connections within auditory-motor circuits change with experience to store memories about self-generated sounds, and how these memories are recalled during subsequent behavior to inform predictions. Together, our experiments demonstrate that the auditory cortex implements precise, movement-based predictions related to self-generated sounds, and provide a tractable inroad towards understanding cortical prediction in general^29,66–71^.

## Methods

### Animals

All experimental protocols were approved by New York University’s Animal Use and Welfare Committee. Male and female wild-type (C57BL/6) mice were purchased from Jackson Laboratories and were subsequently housed and bred in an onsite vivarium. We used 2-4 month old mice for our experiments that were kept on a reverse day-night cycle (12h day, 12 h night).

### Surgeries

For all surgical procedures, mice were anaesthetized under isolfurane (1-2% in O2) and placed in a stereotaxic holder (Kopf), skin was removed over the top of the head, and a Y-shaped titanium headpost (H.E. Parmer) was attached to the skull using a transparent adhesive (Metabond). Mice were treated with an analgesic (Meloxicam SR) and allowed to recover for 5 days prior to training. Following training and 24 to 48 hours prior to electrophysiology, a small craniotomy was made to expose the auditory cortex (~2mm diameter, −2.5mm posterior, 4.2 mm left from bregma). Another small craniotomy was made above the right sensory cortex and a silver-chloride reference electrode was positioned atop the surface of the brain for use as a ground electrode and covered (Metabond). Exposed craniotomies were covered with a silicone elastomer (Kwik-Sil) and the mouse was allowed to recover in its home cage, and an additional training session was performed prior to electrophysiology.

### Behavioral Paradigm Design and Data Collection

We designed a custom head-restrained lever-based behavioral training paradigm where mice push a lever and hear closed-loop sounds. A custom-designed lever (7cm long, 3D printed using Formlabs Form2) was mounted to the post of a rotary encoder (US Digital) 5cm from the lever handle. A magnet (CMS magnetics) was mounted to the bottom of the lever, which was positioned 4cm above a larger static magnet which established the lever resting position and provided light and adjustable movement resistance. The lever handle (top) was positioned adjacent to a tube (Custom, 3D printed using Formlabs Form2) to hold mice and directly below two plate clamps (Altechna) to secure mouse headpost. Lever and mouse apparatus was constructed with Thor-labs components. A water tube, controlled by a solenoid valve (The Lee Company), was positioned in front of the mouse and licking was measured using a custom-mounted (3D printed using Formlabs Form2) IR-beam emitter and receiver. IR signal was titrated and pre-processed using a custom printed circuit board (designed by Melissa Caras and Dan Sanes) to generate a binary TTL signal with IR sensitivity controlled by a potentiometer. Digital signals for licking and lever movement were collected by a data acquisition card (National Instruments) connected to a computer and logged by custom Matlab software (Mathworks, PsychToolBox) and sampled at ~2Khz. Digital inputs of lever movements and licking received sufficient processing in real time to track lick onset and important movement thresholds, which were used to trigger sound events based on user-defined closed-loop rules. Sound output was delivered from the computer to a sound card (RME Fireface UCX), the output of which was routed to an ultrasonic speaker (Tucker Davis Technologies) located lateral to the mouse, ~10cm from the mouse’s right ear. We recorded sounds during test experiments using an ultrasonic microphone to confirm that the lever produced negligible noise and sounds were delivered at a consistent volume. All training was performed in a sound-attenuating booth (Gretch-Ken) to minimize background sound and monitored in real-time via IR video.

We designed two variants of the lever task which differed only in the position used to trigger sounds. In both task variants, water-restricted mice were head-fixed to the behavioral apparatus and presented with the lever and lick-port after ~10 minutes of quiet acclimation. Mice were then allowed to make outwards lever movements at will. For a movement to be considered valid, we required the lever to remain in the home position (~+/− 3mm from rest) for >200 ms prior to initiation. Valid movements that reached a reward threshold (~15mm from home position) elicited a small water reward (5-10uL) when the lever returned to home position. Auditory feedback in the form of a pure tone (50ms duration, ~50dB) was delivered on all trials when the lever crossed a set threshold for the first time in a trial. Mice were trained on a single sound frequency, but the trained frequency was shifted across mice and ranged between 4 and 16 khz. The reward threshold and early sound threshold were slightly different for each mouse depending on their size (+/-2mm), but was consistent across all trials for a mouse after the first few sessions. The only difference between the two task variants was the position at which sound feedback was triggered. For the standard, early-sound variant, sounds were generated when the lever reached 1/3 of the distance between the home position boundary and the reward threshold. For the reward-threshold variant, sounds were triggered using the same threshold that determined reward. To ensure strong coupling between movement and sound, auditory feedback was provided on all trials, regardless of whether mice obeyed the home-position requirement and would subsequently receive a reward.

### Behavioral Training and Data Analysis

Mice received between 19 and 33 sessions of training over roughly two weeks prior to experimentation, with either one or two sessions per day. Prior to the first training session, mice were water-restricted for 24 hours. At the beginning of the first session the reward threshold was set very low (~4mm), reward volume was very high, and a home position time requirement was omitted, enabling the mice to rapidly establish a basic understanding of the task. The reward position was typically extended to at least 80% of the final position by the end of the first hour of training and reached its final position within 3-4 sessions as the mice improved at the task. At this point, the home-time requirement was implemented, and reward threshold remained constant for all remaining sessions. Learning trajectory data in Fig1C only includes sessions after final reward threshold was set. To assist in training and further differentiate the tone relevance for the two tasks, mice trained with a reward-threshold tone received auditory feedback from the onset of the first session, but mice trained with an early-position tone did not hear auditory feedback until the final reward threshold was established.

Lever trajectories and licking were analyzed post-hoc to assess mouse behavioral patterns. For each training session, as well as the electrophysiology behavior session, we measured the average lever trajectory of all movements that reached the sound threshold, normalized to the distance of the reward threshold. For each mouse, we also measured the average maximum lever position and movement duration across lever pushes and created a histogram of the maximum lever position of all individual movements. The empirical coupling between movement, sound, and reward was averaged across the last three training sessions for each animal, measured as the fraction of movements that generated a sound and the fraction of sounds that were followed by a reward.

To assess learned expectations for self-generated sounds, we introduced tone deviations on a subset of trials and measured changes in these lever properties. Omissions of expected sounds occurred on the training session prior to craniotomy, when sounds were omitted on 1 in 15 trials randomly interleaved, beginning after mice had performed at least 30 standard trials. For mice trained to expect a reward-threshold tone we also performed a second, separate behavior session where tones were played slightly earlier than expected (15%, ~3mm) on 1 in 15 trials. Deviants were presented at a low rate to probe behavior without disrupting learned expectations prior to electrophysiological recording. Behavioral deviations to frequency probes, where sound was played at the correct lever position but shifted one octave above or below the expected tone, were measured on electrophysiology day at a rate of 1 to 1. Deviant auditory feedback did not alter the requirements for reward. For each probe type, we measured the average lever trajectory over time, average movement duration, average maximum lever position, average lick onset time, and lick rate (200ms centered around movement end) for the first 30 probe trials and compared these values to the same measurements calculated for all interleaved standard trials occurring between the first and last probe trial. Only trials that reached the sound-threshold were included in analysis, except for during the early-tone probe session, where movement inclusion for standard training was based on the distance threshold of the early tone to enable a fair comparison of movement distances. We also measured the root-mean-square-error between average lever trajectories for interleaved standard and probe trials, as well as between the 30 trials directly preceding deviant tone onset and the 30 trials directly after deviant onset regardless of trial type. We compared each RMSE value to the RMSE difference of average trajectories from the same trial pool but with shuffled trial identities. The shuffled RMSE value reported here is the average of 1000 iterations of shuffled RMSE difference calculation.

### Electrophysiological Recording and Analysis

Following training, behavioral probe experiments, surgical opening of a craniotomy, and one subsequent training session, mice were positioned in the behavioral apparatus and a 128-channel electrode (128AxN, Masmanidis Lab) was lowered into the auditory cortex orthogonal to the pial surface^39^. The electrode was connected to a digitizing head stage (Intan) and electrode signals were acquired at 30khz, monitored in real time, and stored for offline analysis (OpenEphys). The probe was allowed to settle for at least 20 minutes, at which point the lever and lick-port were introduced and mice were allowed to make lever movements at will as in any other training session. After performing at least 30 standard lever movements, we unexpectedly began a probe session in which mice heard either the lever-associated sound or a single probe sound one octave away (same duration and intensity) on 50% of pseudorandom trials shuffled in batches of 30 tones. Disbursements of rewards was blind to tone identity. The standard frequency-probe session was followed by a broader frequency-probe session where lever pushes generated tones randomly at half octave intervals around the lever-associated tone. Following probe sessions, the lever was removed, and we played the same sounds to the animal in a passive context.

After recording, electrical signals were processed and the action-potentials of individual neurons were sorted using Kilosort2, and manually reviewed in Phy2 based on reported contamination, waveform principal component analysis, and inter-spike interval histograms. Units were only included in analysis if they maintained their baseline firing rate for the duration of the experiment (+/− 30%), and demonstrated a non-fast-spiking waveform, separated by plotting peak to valley ratio against action potential width (ms). Tone-evoked action potential responses were measured in 5ms bins and aligned to sound onset for each neuron for each tone type. Heat maps of individual neuronal firing (Fig 1G and Fig 2E) show raw firing rates averaged across all trials with a threshold at 0 and 30hz for all recorded neurons. In order to study the neural representations of sounds in the cortex, subsequent analysis of sound-responses includes all neurons that were responsive (p<0.1) to a given tone in either the active or passive condition measured as an increase in firing rate from baseline (average over 100ms prior to stimulus) during the sound response window (10-60ms post stimulus) across trials. We used this inclusion criteria because we wanted to capture a broad range of neurons likely to be involved in processing cortical sounds which need not be limited to classically strongly sound-responsive neurons.^72^ Tone-evoked responses were visualized by calculating the average firing rate in 5ms bins over time and subtracting baseline activity (average firing rate in the 100ms window prior to sound onset). Tone-evoked responses were quantified by comparing the driven firing rate (10-60ms after sound onset) to the baseline firing rate (100ms window prior to sound onset). To measure the movement-based modulation of each neuron’s responses to the lever-associated or probe tone, we compared the neural sound response in our analysis window to the same sound in the active and passive condition using a radial modulation index. Radial modulation was calculated as the theta value resulting from a cartesian to polar transformation of the response strength in the active condition compared to the response strength in the passive condition. Theta values were converted to a scale of +/− 2 and rotated such that a value of 0 corresponded to equal responses across the two conditions. These neuronal modulation values are calculated and displayed just for the pool of neurons responsive to a given sound, and were averaged across all cells, or within an animal. We followed a similar procedure for each probe tones when there was more than one probe frequency in a given experiment. To assess the stability of neural responses to a sound throughout an entire experiment, we measured the average population response of all sound-responsive neurons on each trial. We then measured the average modulation for quartiles of trials for the expected tone during standard training and independently for both the expected and unexpected sound during the probe section. The above analyses were performed in parallel for separate pools of mice trained on the two task variants.

### Experimental Variants and Specific Analyses

In a subset of animals, we performed electrophysiological recording of mice that had been trained on an identical version of the lever task but without sound feedback (7-10 sessions). On experiment day mice first performed silent lever pushes for 20-50 trials, then we performed a broad frequency probe (see above), followed by a session of standard training with a single tone heard early in every movement, followed by a post-training broad frequency probe. These data were used to assess both movement responses and sound responses prior to movement-sound coupling.

In the experimental variant focused on background sounds, mice received identical training on the early-tone task variant, but the normal frequency-probe experiment was accompanied by the delivery of the lever-associated and probe tone played randomly using intervals between 0.5 and 1.5s in the background. Background sounds were also introduced on the 5 sessions prior to electrophysiology to acclimate the mice to this disruptive experience. The probe session was followed by lever removal and passive replay of the two sounds. Analysis of lever-generated tones was performed as described above for standard frequency-probe experiments. Neural responses to background tones were analyzed for each frequency if they fell within 0 to 200ms, or 200 to 400ms, or 400 to 600ms prior to lever movement onset, and if there was no licking or lever movements in the 500ms prior to the tone. Neural responses were compared to passive tones to generate a modulation value. In the experimental variant focused on time deviants within the lever movement, mice were again trained on the standard early-tone behavior prior to electrophysiology. During recording, mice experience the expected frequency and a probe frequency at three different positions that were identical within and across mice: the expected position, the reward threshold, and halfway through the return phase. 90% of tones occurred at the expected position and frequency, and with the remaining 10% distributed across the other 5 position and frequency combinations. Reward delivery was unaffected by changes in sound position, and a tone only played on the return phase if a given movement reached the reward threshold. This created a small number of omission trials which were used to assess movement signals in the absence of sound. Neural responses to lever sounds. Neural responses to sounds at different time points were compared to passively heard tones at the end of the experiments, with neurons contributing to modulation values if they responded (p<0.1) to that tone type or the same frequency heard in the passive condition.

In the experimental variant focused on movement specificity, we delivered sounds during licking in a subset of animals only during electrophysiology experiments. Directly following the broad lever-initiated frequency-probe, we randomly delivered the same set of tones at half-octave intervals but triggered by individual mouse licks that occurred naturally during behavior, especially in the seconds right after movement termination. During the session of licking-initiated sounds, the lever continued to generate random half-octave interval tones. We performed a normal calculation of neural sound responses and modulation compared to passively heard tones for licking-evoked sounds that occurred outside of lever movement.

Laminar assignment of neural identity was performed for all experiments based on consistent features of our observed current source density (example in Fig5A). Current source density analysis following passively heard tones at the end of each experiment was performed independently for each probe shank, with each shank treated as a composite of three different linear arrays^73^. Each feature was assigned a consistent neural depth across animals to generate a consistent depth and laminar scale across animals. The mapping of CSD features to numerical depths and putative layers is an estimate based on our observed empirical depths, CSD analysis in other experiments, and studies of laminar architecture in the auditory cortex^74–76^. While numerical depths are inferred, neurons are correctly ordered with respect to one another and to key CSD features. For averages of neural features across depth, we performed a moving average across 50um centered on each cortical depth (5um bins).

Neural representations of self-generated sounds were determined by measuring the responsiveness of each recorded neuron to the expected and unexpected tone in both the active and passive condition (p<0.05). We categorized neurons as responsive to both sounds in all conditions, both sounds but only during movement, specific to a frequency regardless of behavioral context, or specific to a given tone only during movement. Importantly, the first two groups of neurons (all sounds, movement sounds) are the same population of neurons across both tone types, but frequency-driven neurons and neurons responsive specifically to self-generated sounds are different groups of neurons for the expected sound and unexpected sound.

Movement responses in the auditory cortex were measured on omission trials (see above) that reached at least 66% of the distance to reward threshold. A broad pool of movement-activated neurons was identified by measuring responsiveness (p<0.01) in the 100ms before or after each of 3 positions during movement (15%, 50%, 80% of reward threshold) on both the outbound and return phase of movements, using a consistent baseline 400-500ms prior to movement onset. Movement signals for all responsive neurons were visualized relative to the time of expected sound position in all figures, except in Fig 4C,D where average neural firing is shown aligned to each of the six time points used to identify responsive neurons. For each of these neurons, we z-scored average neural firing during lever movements based on the average and standard-deviation of neural firing rate across trials in the baseline window (500 to 400ms prior to sound). If a neuron reached a threshold of 2.5 standard deviations above baseline, we calculated a time of peak firing as well as an onset time measured as the crossing of 0.5 standard deviations prior to the peak. We also measured the fraction of neurons that were sound-responsive at each cortical depth by comparing the number of responsive neurons in a 50um window around each cortical depth (5um bins) compared to the total count of neurons in the same window. Movement responses for the core dataset (early-tone training and frequency-deviant probes) were calculated for the 100ms prior to sound onset compared to the same baseline window (400-500ms prior to sound).

### Clustering Analysis

For unbiased clustering analysis we logged parameters characterizing the activity of each neuron in our core dataset to each of our behavioral and sound variables: Expected Frequency sound heard actively and passively, probe frequency-sound heard actively and passively, and movement signals in the time prior to sound onset. Neural response to each variable was measured both as z-scored neural activity level, and as a baseline subtracted response rate (Sound response: 0-60ms vs 100ms prior; Movement: 100ms prior to sound onset vs 400-500ms prior). Each of the 10 dimensions was pre-processed prior to clustering, comprised of log normalization, thresholding of outliers to a boundary value, and then centering the distribution between −1 and 1. We then performed k-medoids clustering^45^ on the processed dataset, a variant of k-means clustering that uses individual datapoints as group centers and is more robust to outliers. We performed clustering analysis for 3-30 clusters, performing the elbow test and silhouette test to measure the quality of clustering and determine the appropriate number of clusters (7 for core dataset). To insure that the functional properties of our groups were not driven by cluster number, we looked qualitatively at cluster output groups at a variety of k values and found similar functional populations, albeit split or merged across different numbers of groups. Similarly, we performed clustering after using an alternate pre-processing approach where neuron values within each dimension were ordered and then fit to a normal distribution, which yielded similar clustering results. Cluster results were mapped onto a 2-dimensional representation of the data generated using UMAP for visualization^77^.

Neurons were sorted by cluster, and by average activity within cluster, and visualized using Z-scored neural activity over time to each stimulus type and probability response values to each variable. We then characterized select groups by plotting PSTHs, response graphs, modulation graphs, and laminar plots including all neurons in a given group. Response graphs reflect the baseline-subtracted responses that went into clustering, modulation graph includes only neurons that are response to a given tone in either the active or passive condition, and depth plots show both the number of cells by depth and the fraction of cells in each layer.

We additionally clustered neural activity in the dataset that included omission probes in addition to frequency probes. Clustering was performed with identical dimensions and pre-processing, except that movement activity was measured at two time points: prior to the time of expected sound, and just after the time of expected sound (0-60ms post expected onset vs 100ms prior). Clustering was robust to small differences in cluster number and pre-processing, and cluster analysis determined a k value of 10 groups. Results were visualized similarly but included the additional movement time point following the time of expected auditory feedback.

### Statistical Analysis

Throughout, animal values are denoted by a capital N while cell values are denoted by a lowercase n. Unless otherwise reported, all error bars denote standard deviation, and all statistical comparisons were performed using a two sided, non-paired, non-parametric rank sum test. Additionally, all neural percentages reported use a denominator of all neurons, or all neurons in a given depth rather than a smaller denominator comprising just sound-responsive neurons. For broad frequency-probe experiments, modulation of neurons to the expected tone were compared to modulation of neurons to all probe tones pooled together. For statistical analysis of RMSE differences for interleaved and pre vs post comparisons, we performed a paired Wilcoxon test between the empirical RMSE values for each condition compared to the average of 1000 RMSE values generated by random trial assignment. Statistical comparison of laminar distributions were performed using a Kolmogorov-Smirnov test, and distributions of movement response properties were compared using both the Kolmogorov-Smirnov test and the f-test. The relationship between movement signal strength and timing was performed using linear regression and correlation coefficient analysis with all p and R values reported.

## Acknowledgements

We thank Dr. Dan Sanes, Dr. Adam Carter, Dr. Alex Williams, Dr. Alessandro La Chioma, and Ralph E. Peterson for their thoughtful and valuable comments on this manuscript. We thank members of the Schneider lab for fruitful discussions regarding experimental design, data analysis, and interpretation. We thank Hoda Ansari and Jessica A. Guevara for their expert animal care and technical support. This research was supported by the National Institutes of Health (T32-MH019524 to N.A., 1R01-DC018802 to D.M.S.); a Career Award at the Scientific Interface from the Burroughs Wellcome Fund (D.M.S); fellowships from the Searle Scholars Program, the Alfred P. Sloan Foundation, and the McKnight Foundation (D.M.S.); and an investigator award from the New York Stem Cell Foundation (D.M.S). D.M.S. is a New York Stem Cell Foundation - Robertson Neuroscience Investigator.

## Author contributions

Conceptualization, N.J.A., and D.M.S.; Investigation, N.J.A. and W.Z..; Writing – Original Draft, N.J.A. and D.M.S.; Writing – Review & Editing, N.J.A., W.Z., and D.M.S. Supervision, D.M.S.; Funding Acquisition, D.M.S.

## Data and Code Availability Statements

The datasets generated during and/or analyzed during the current study are available from the corresponding author on reasonable request.

All code used in the collection and analysis of data during the current study are available from the corresponding author on reasonable request.

## References

1. Wolpert, D. M., Ghahramani, Z. & Jordan, M. I. An internal model for sensorimotor integration. Science (80.-). 269, 1880–1882 (1995).

2. Nelson, A. et al. A circuit for motor cortical modulation of auditory cortical activity. J. Neurosci. 33, 14342–14353 (2013).

3. Keller, G. B. & Mrsic-Flogel, T. D. Predictive Processing: A Canonical Cortical Computation. Neuron 100, 424–435 (2018).

4. Schneider, D. M., Nelson, A. & Mooney, R. A synaptic and circuit basis for corollary discharge in the auditory cortex. Nature 513, 189–194 (2014).

5. Schneider, D. M. & Mooney, R. How movement modulates hearing. Annu. Rev. Neurosci. 41, 553–572 (2018).

6. Singla, S., Dempsey, C., Warren, R., Enikolopov, A. G. & Sawtell, N. B. A cerebellum-like circuit in the auditory system cancels responses to self-generated sounds. Nat. Neurosci. 20, 943–950 (2017).

7. Clancy, K. B., Orsolic, I. & Mrsic-Flogel, T. D. Locomotion-dependent remapping of distributed cortical networks. Nat. Neurosci. 22, 778–786 (2019).

8. Niell, C. M. & Stryker, M. P. Modulation of Visual Responses by Behavioral State in Mouse Visual Cortex. Neuron 65, 472–479 (2010).

9. Ayaz, A. et al. Layer-specific integration of locomotion and sensory information in mouse barrel cortex. Nat. Commun. 10, (2019).

10. Ayaz, A., Saleem, A. B., Schölvinck, M. L. & Carandini, M. Locomotion controls spatial integration in mouse visual cortex. Curr. Biol. 23, 890–894 (2013).

11. Brainard, M. S. & Doupe, A. J. Auditory feedback in learning and maintenance of vocal behaviour. Nat. Rev. Neurosci. 1, 31–40 (2000).

12. Rao, R. P. N. & Ballard, D. H. Predictive coding in the visual cortex: a functional interpretation of some extra-classical receptive-field effects. Nat. Neurosci. 2, 79–87 (1999).

13. Hickok, G., Houde, J. & Rong, F. Sensorimotor Integration in Speech Processing: Computational Basis and Neural Organization. Neuron 69, 407–422 (2011).

14. Bastos, A. M. et al. Canonical Microcircuits for Predictive Coding. Neuron 76, 695–711 (2012).

15. Steinmetz, N. A., Zatka-Haas, P., Carandini, M. & Harris, K. D. Distributed coding of choice, action and engagement across the mouse brain. Nature 576, 266–273 (2019).

16. Stringer, C. et al. Spontaneous behaviors drive multidimensional, brainwide activity. Science (80-.). 364, (2019).

17. Musall, S., Kaufman, M. T., Juavinett, A. L., Gluf, S. & Churchland, A. K. Single-trial neural dynamics are dominated by richly varied movements. Nat. Neurosci. 22, 1677–1686 (2019).

18. McGinley, M. J. et al. Waking State: Rapid Variations Modulate Neural and Behavioral Responses. Neuron 87, 1143–1161 (2015).

19. Hasenstaub, A., Sachdev, R. N. S. & McCormick, D. A. State changes rapidly modulate cortical neuronal responsiveness. J. Neurosci. 27, 9607–9622 (2007).

20. Bennett, C., Arroyo, S. & Hestrin, S. Subthreshold mechanisms underlying state-dependent modulation of visual responses. Neuron 80, 350–357 (2013).

21. Kuchibhotla, K. V. et al. Parallel processing by cortical inhibition enables context-dependent behavior. Nat. Neurosci. 20, 62–71 (2017).

22. Zhou, M. et al. Scaling down of balanced excitation and inhibition by active behavioral states in auditory cortex. Nat. Neurosci. 17, 841–850 (2014).

23. Polack, P.-O., Friedman, J. & Golshani, P. Cellular mechanisms of brain state-dependent gain modulation in visual cortex. 16, (2013).

24. Leinweber, M., Ward, D. R., Sobczak, J. M., Attinger, A. & Keller, G. B. A Sensorimotor Circuit in Mouse Cortex for Visual Flow Predictions. Neuron 95, 1420–1432.e5 (2017).

25. Zagha, E., Casale, A. E., Sachdev, R. N. S., McGinley, M. J. & McCormick, D. A. Motor cortex feedback influences sensory processing by modulating network state. Neuron 79, 567–578 (2013).

26. Lee, S., Kruglikov, I., Huang, J., Fishell, G. & Rudy, B. A disinhibitory circuit mediates motor integration in the somatosensory cortex. Nat. Publ. Gr. (2013) doi:10.1038/nn.3544.

27. Reznik, D., Guttman, N., Buaron, B., Zion-Golumbic, E. & Mukamel, R. Action-locked Neural Responses in Auditory Cortex to Self-generated Sounds. Cereb. Cortex 31, 5560–5569 (2021).

28. Schneider, D. M., Sundararajan, J. & Mooney, R. A cortical filter that learns to suppress the acoustic consequences of movement. Nature 561, 391–395 (2018).

29. Rummell, B. P., Klee, J. L. & Sigurdsson, T. Attenuation of responses to self-generated sounds in auditory cortical neurons. J. Neurosci. 36, 12010–12026 (2016).

30. Eliades, S. J. & Wang, X. Neural substrates of vocalization feedback monitoring in primate auditory cortex. Nature 453, 1102–1106 (2008).

31. Flinker, A. et al. Single-trial speech suppression of auditory cortex activity in humans. J. Neurosci. 30, 16643–16650 (2010).

32. Knolle, F., Schwartze, M., Schröger, E. & Kotz, S. A. Auditory Predictions and Prediction Errors in Response to Self-Initiated Vowels. Front. Neurosci. 13, 1–11 (2019).

33. Zmarz, P. & Keller, G. B. Mismatch Receptive Fields in Mouse Visual Cortex. Neuron 92, 766–772 (2016).

34. Jordan, R. & Keller, G. B. Opposing Influence of Top-down and Bottom-up Input on Excitatory Layer 2/3 Neurons in Mouse Primary Visual Cortex. Neuron 108, 1194–1206.e5 (2020).

35. Keller, G. B., Bonhoeffer, T. & Hübener, M. Sensorimotor Mismatch Signals in Primary Visual Cortex of the Behaving Mouse. Neuron 74, 809–815 (2012).

36. Muzzu, T. & Saleem, A. B. Feature selectivity can explain mismatch signals in mouse visual cortex Graphical abstract. CellReports 37, 109772 (2021).

37. Natan, R. G. et al. Complementary control of sensory adaptation by two types of cortical interneurons. Elife 4, 1–27 (2015).

38. Wiltschko, A. B. et al. Mapping Sub-Second Structure in Mouse Behavior. Neuron 88, 1121–1135 (2015).

39. Yang, L., Lee, K., Villagracia, J. & Masmanidis, S. C. Open source silicon microprobes for high throughput neural recording. J. Neural Eng. 17, (2020).

40. Bao, S., Chan, V. T. & Merzenich, M. M. Cortical remodelling induced by activity of ventral tegmental dopamine neurons. Nature 412, 79–83 (2001).

41. Fritz, J., Shamma, S., Elhilali, M. & Klein, D. Rapid task-related plasticity of spectrotemporal receptive fields in primary auditory cortex. (2003) doi:10.1038/nn1141.

42. Cynx, J. & Von Rad, U. Immediate and transitory effects of delayed auditory feedback on bird song production. Anim. Behav. 62, 305–312 (2001).

43. Stuart, A., Kalinowski, J. & Rastatter, M. P. Effect of delayed auditory feedback on normal speakers at two speech rates. Cit. J. Acoust. Soc. Am. 111, 2237 (2002).

44. Clayton, K. K. et al. Auditory Corticothalamic Neurons Are Recruited by Motor Preparatory Inputs. Curr. Biol. 31, 310–321.e5 (2021).

45. Park, H. S. & Jun, C. H. A simple and fast algorithm for K-medoids clustering. Expert Syst. Appl. 36, 3336–3341 (2009).

46. Angeloni, C. & Geffen, M. N. Contextual modulation of sound processing in the auditory cortex. Curr. Opin. Neurobiol. 49, 8–15 (2018).

47. Goard, M. & Dan, Y. Basal forebrain activation enhances cortical coding of natural scenes. Nat. Neurosci. 12, 1444–1449 (2009).

48. Nelson, A. & Mooney, R. The Basal Forebrain and Motor Cortex Provide Convergent yet Distinct Movement-Related Inputs to the Auditory Cortex. Neuron 90, 635–648 (2016).

49. Roth, M. M. et al. Thalamic nuclei convey diverse contextual information to layer 1 of visual cortex. Nat. Neurosci. 19, 299–307 (2016).

50. Ji, X. Y. et al. Thalamocortical Innervation Pattern in Mouse Auditory and Visual Cortex: Laminar and Cell-Type Specificity. Cereb. Cortex 26, 2612–2625 (2016).

51. Audette, N. J., Urban-Ciecko, J., Matsushita, M. & Barth, A. L. POm Thalamocortical Input Drives Layer-Specific Microcircuits in Somatosensory Cortex. Cereb. Cortex 28, 1312–1328 (2018).

52. Sawtell, N. B. & Williams, A. Transformations of electrosensory encoding associated with an adaptive filter. J. Neurosci. 28, 1598–1612 (2008).

53. Kloc, M. & Maffei, A. Target-specific properties of thalamocortical synapses onto layer 4 of mouse primary visual cortex. J. Neurosci. 34, 15455–15465 (2014).

54. Constantinople, C. M. & Bruno, R. M. Deep Cortical Layers are Activated Directly by Thalamus. Science (80-.). 1591, 1591–1594 (2013).

55. Crandall, S. R., Patrick, S. L., Cruikshank, S. J. & Connors, B. W. Infrabarrels Are Layer 6 Circuit Modules in the Barrel Cortex that Link Long-Range Inputs and Outputs. Cell Rep. 21, 3065–3078 (2017).

56. Viaene, A. N., Petrof, I. & Sherman, S. M. Synaptic properties of thalamic input to layers 2/3 and 4 of primary somatosensory and auditory cortices. J. Neurophysiol. 105, 279–292 (2011).

57. Viaene, A. N., Petrof, I. & Murray Sherman, S. Synaptic properties of thalamic input to the subgranular layers of primary somatosensory and auditory cortices in the mouse. J. Neurosci. 31, 12738–12747 (2011).

58. Sermet, B. S. et al. Pathway-, layer-and cell-type-specific thalamic input to mouse barrel cortex. Elife 8, 1–28 (2019).

59. Eliades, S. J. & Tsunada, J. Auditory cortical activity drives feedback-dependent vocal control in marmosets. doi:10.1038/s41467-018-04961-8.

60. Makino, H., Hwang, E. J., Hedrick, N. G. & Komiyama, T. Circuit Mechanisms of Sensorimotor Learning. Neuron 92, 705–721 (2016).

61. Barth, A. L. & Poulet, J. F. A. Experimental evidence for sparse firing in the neocortex. Trends Neurosci. 35, 345–355 (2012).

62. Crochet, S. & Petersen, C. C. H. Correlating whisker behavior with membrane potential in barrel cortex of awake mice. Nat. Neurosci. 9, 608–610 (2006).

63. Haider, B. et al. Synaptic and Network Mechanisms of Sparse and Reliable Visual Cortical Activity during Nonclassical Receptive Field Stimulation. Neuron 65, 107–121 (2010).

64. Vinje, W. E. & Gallant, J. L. Sparse coding and decorrelation in primary visual cortex during natural vision. Science (80-.). 287, 1273–1276 (2000).

65. Francis, N. A. et al. Small Networks Encode Decision-Making in Primary Auditory Cortex. Neuron 97, 885–897.e6 (2018).

66. Cooke, S. F., Komorowski, R. W., Kaplan, E. S., Gavornik, J. P. & Bear, M. F. Visual recognition memory, manifested as long-term habituation, requires synaptic plasticity in V1. 2, (2015).

67. Demarchi, G., Sanchez, G. & Weisz, N. Automatic and feature-specific prediction-related neural activity in the human auditory system. Nat. Commun. 10, (2019).

68. Gavornik, J. P. & Bear, M. F. Learned spatiotemporal sequence recognition and prediction in primary visual cortex. Nat. Neurosci. 17, 732–737 (2014).

69. Kato, H. K., Gillet, S. N. & Isaacson, J. S. Flexible Sensory Representations in Auditory Cortex Driven by Behavioral Relevance. Neuron 88, 1027–1039 (2015).

70. Voigts, J., Deister, C. A. & Moore, C. I. Layer ensembles can selectively regulate the behavioral impact and layer-specific representation of sensory deviants. Elife 9, 1–39 (2020).

71. Garner, A. R. & Keller, G. B. A cortical circuit for audio-visual predictions. bioRxiv 2020.11.15.383471 (2020) doi:10.1038/s41593-021-00974-7.

72. Insanally, M. N. et al. Spike-timing-dependent ensemble encoding by non-classically responsive cortical neurons. Elife 8, 1–31 (2019).

73. Pettersen, K. H., Devor, A., Ulbert, I., Dale, A. M. & Einevoll, G. T. Current-source density estimation based on inversion of electrostatic forward solution: Effects of finite extent of neuronal activity and conductivity discontinuities. J. Neurosci. Methods 154, 116–133 (2006).

74. Chang, M. & Kawai, H. D. A characterization of laminar architecture in mouse primary auditory cortex. Brain Struct. Funct. 223, 4187–4209 (2018).

75. Szymanski, F. D., Rabinowitz, N. C., Magri, C., Panzeri, S. & Schnupp, J. W. H. The laminar and temporal structure of stimulus information in the phase of field potentials of auditory cortex. J. Neurosci. 31, 15787–15801 (2011).

76. Muramatsu, S., Toda, M., Nishikawa, J. & Tateno, T. Sound- and current-driven laminar profiles and their application method mimicking acoustic responses in the mouse auditory cortex in vivo. Brain Res. 1721, 146312 (2019).

77. McInnes, L., Healy, J. & Melville, J. UMAP: Uniform Manifold Approximation and Projection for Dimension Reduction. bioRxiv (2018).

