## Supplemental Figures for "Temporally precise movement-based predictions in the mouse auditory cortex"

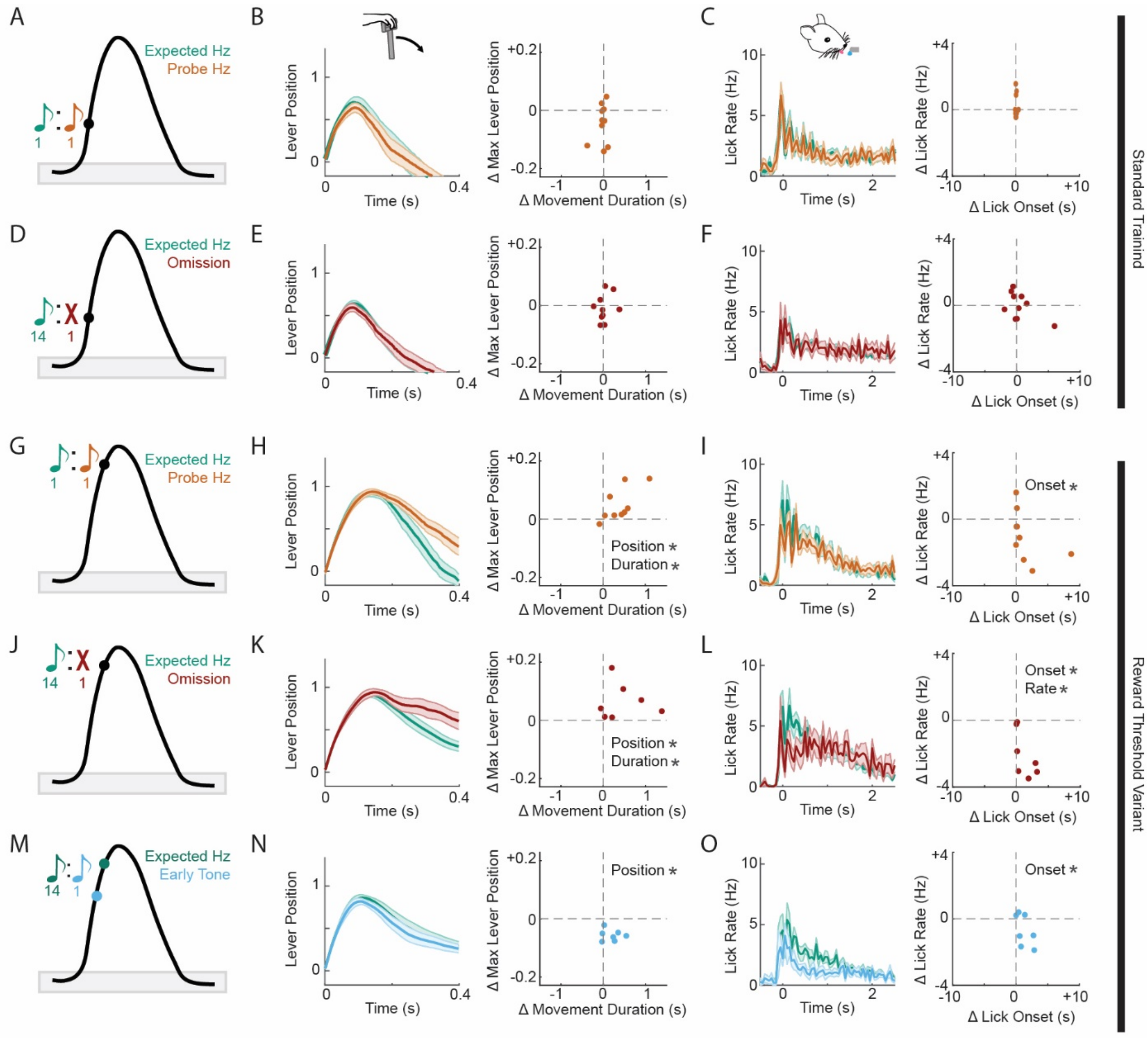


**Figure S1. Self-generated sounds are used to guide ongoing behavior when they are expected at the reward threshold, but not early in movement.** (A) Schematic of probe experiment where the expected, early tone is shifted one octave in a consistent direction on 50% of randomly interleaved trials. (B) Global average lever trajectory +/- standard error across mice for interleaved expected-sound trials and probe trials (left), measured for all movements that generated a sound, aligned to movement onset. Difference in maximum lever reach and movement duration for probe trials compared to interleaved expected-sound trials. (C) Global average rate of mouse licking aligned to time of lever-return (left), and difference in licking rate and licking onset for frequency probe trials compared to interleaved control trials (right). (D-F) Same as (A-C), but for sound omission deviant experiment where the expected, early tone is omitted on 1 in 15 randomly interleaved trials. (G-L) Same as (A-F), but with sounds frequency shifted (G-I) or omitted (J-L) in an identical fashion but for mice trained to expect the self-generated sound at the reward threshold. (M-O) Same as (J-L), but with tones played 15% early on 1 in 15 trials randomly interleaved with expected tones occurring at the reward threshold. *$p < 0.05$

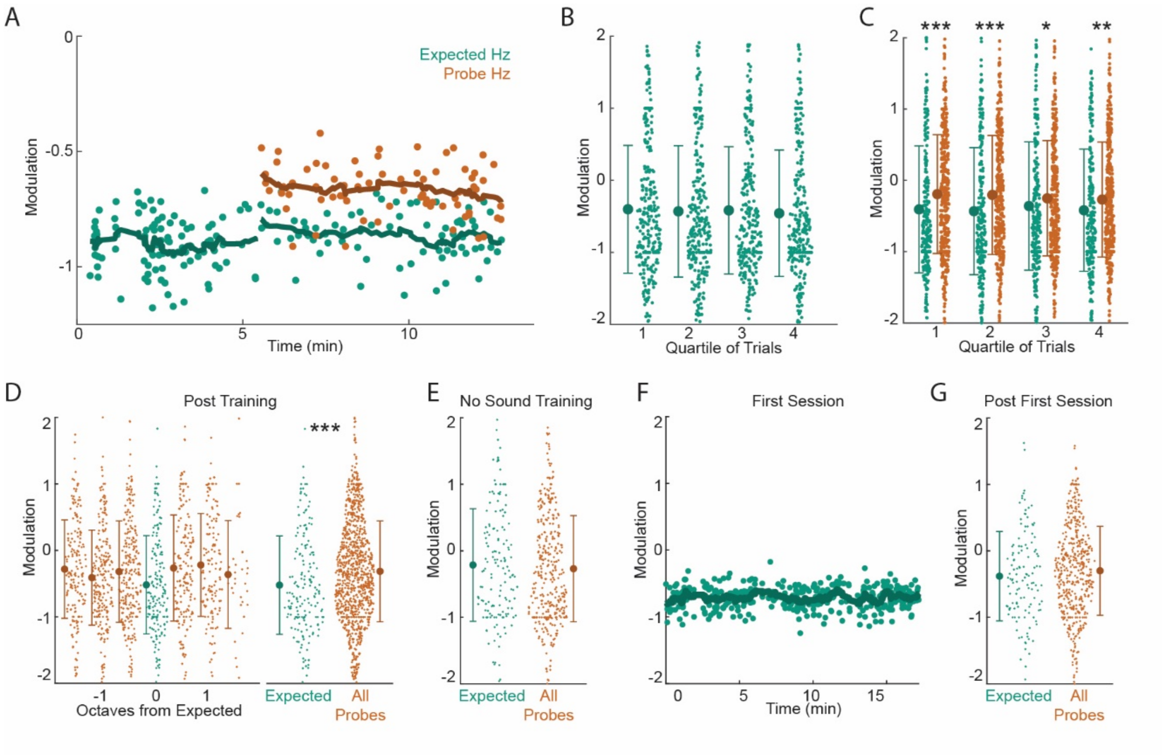


**Figure S2. Expected, self-generated sounds are subject to a notch filter that is learned, and stable over time.** (A) Average modulation across the neural population for individual trials over time in an example animal, with across-trial moving mean. Experiment and data begin with a section of standard training followed by a frequency probe section. (B, C) Average neural modulation (N=10) for trial quartiles of the standard behavior section (B) and the frequency-probe section (C). (D) Individual neuron modulation and average modulation of frequencies delivered at half-octave intervals, organized based on their distance in octaves from the expected sound (left). Statistical comparison (right) of modulation of the expected tone compared to neural response modulation pooled across all other tones. (E) Comparison of modulation for the expected sound and unexpected frequency sounds delivered prior to the first movement-sound training day in mice trained to push the lever in the absence of sound. Note, expected sound is designated as the frequency that mice will subsequently be trained on. (F) Average modulation across the neural population for individual trials over time on the first movement-sound training day in mice trained to complete the lever task in the absence of sound. Individual mouse example. (G) Comparison of modulation for the expected sound and unexpected frequency sounds delivered after the fist movement-sound training session in mice trained previously in the absence of sound. Note that data in (E-G) are collected in a single training session for each animal (N = 4). * $p < 0.05$ **$p < 0.005$ ***$p<0.0005$

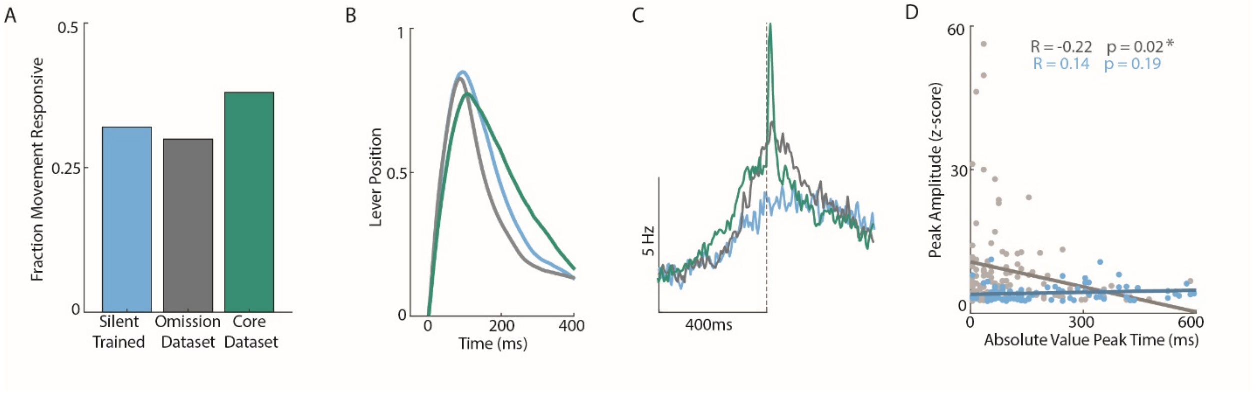


**Figure S4**. **Movement trajectories and movement-responsive neurons across datasets.** (A) Fraction of neurons with movement responses for silent trained dataset (blue), omission dataset (gray), and core dataset (cyan). Note, core dataset can only measure movement responses that occur prior to sound onset. (B) Average lever trajectory over time across all mice for each condition. All movements that crossed the sound threshold early in movement are included. (C) Average firing rate of movement responsive neurons in auditory cortex for each group, aligned to the time of sound (core dataset) or expected sound on omission trials (silent-trained and omission dataset). (D) Plot of movement response amplitude based on the time of movement peak measured as absolute time difference from time of expected sound, with linear regression plotted. R value and p value reported for omission dataset (gray) and silent trained dataset (blue).

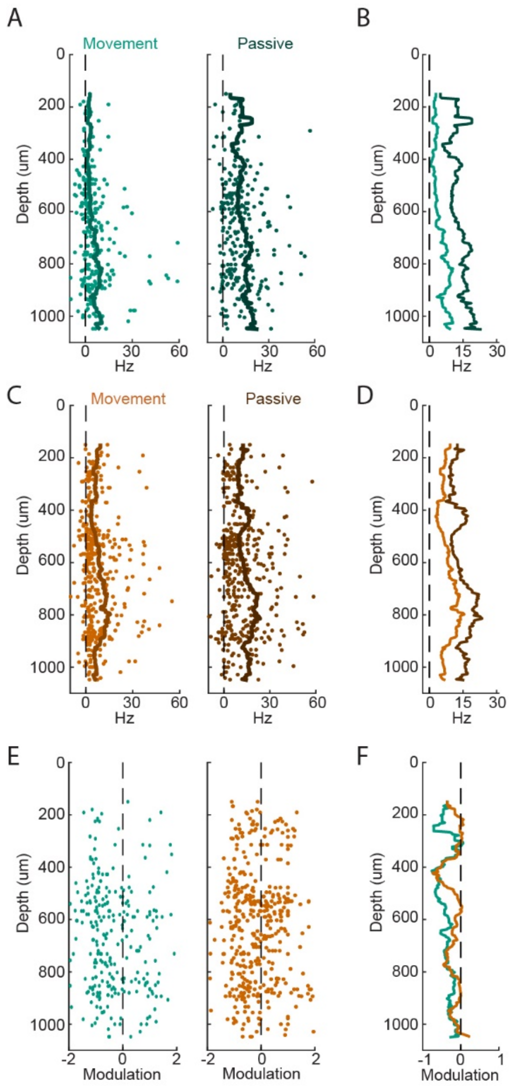


**Figure S5. Detailed sound responses across relative cortical depths.** (A) Average individual neuron responses to the lever-associated sound heard in the active (light,left) or passive (dark,right) context plotted by neural depth. Moving average computed across 50um. (B) Overlay of moving average for active and passive tones on consistent scale. (C,D) Same as A,B but for the unexpected, probe tone. (E) Average neural modulation for the expected sound (left) or probe sound (right) plotted by neural depth. (F) Moving average of modulation of expected and probe sounds by depth.

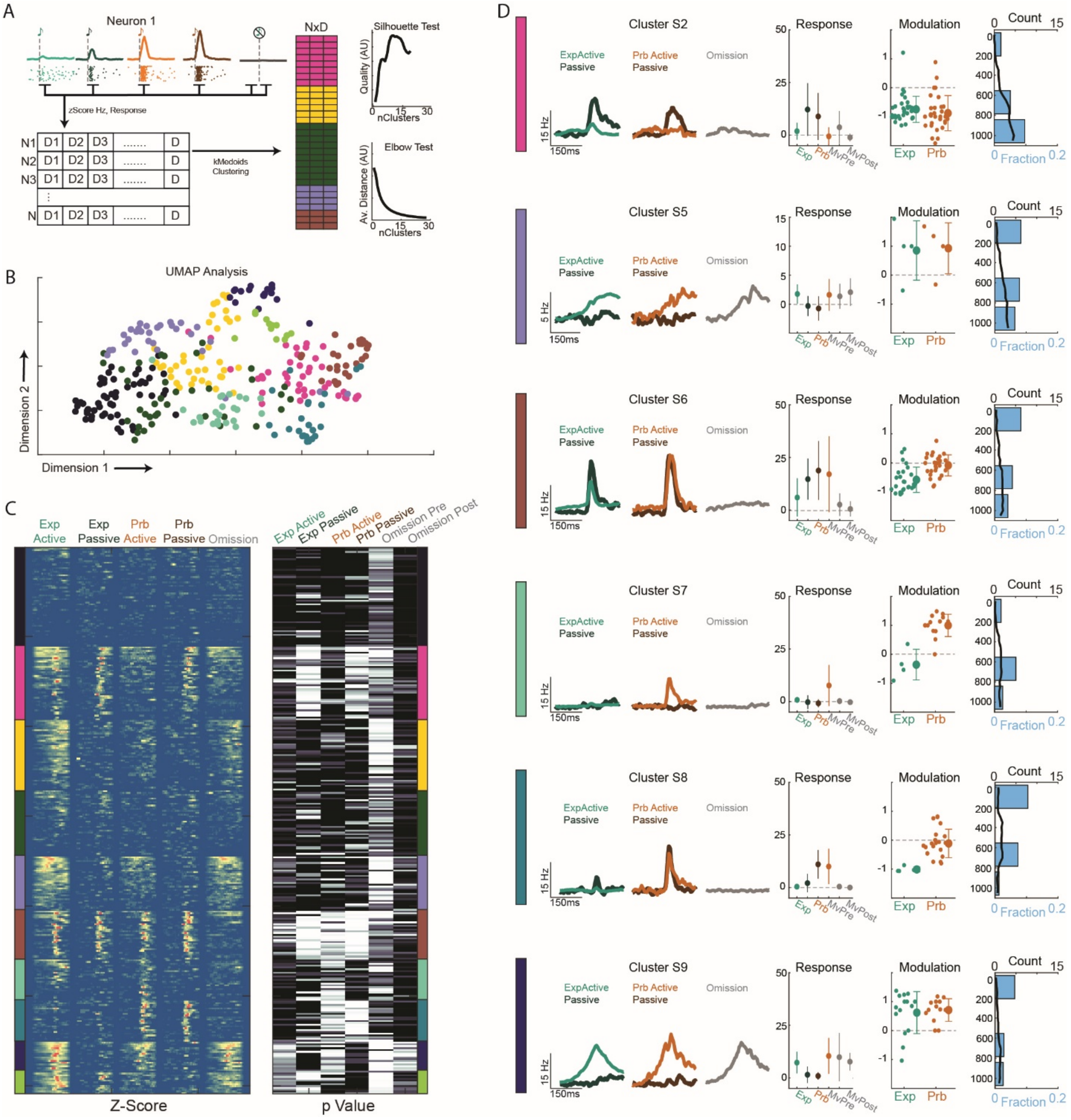

**Figure S7. Measuring neural responses to both frequency and omission deviations improves clustering resolution of movement-responsive functional groups.** (A) Schematic (left) of clustering process, where individual neurons are characterized by their responses to different task variables and then grouped using k-medoids clustering based on these dimensions (Omission dataset, N5, see methods). Silhouette test (top right) showing the quality of clustering for different cluster numbers (k-values), and elbow test (bottom right) showing the average distance of all data points to group center as a function of cluster number. (B) Visualization of cluster groups (k = 10) by plotting each neuron in a low-dimension projection of our feature-space generated by UMAP and color coded by cluster identity. (C) Neuron response patterns sorted by cluster group (designated by color), shown as the z-scored activity pattern over time to each sound type, and as response p-values following sound onset and omission (0-60ms post sound time vs 100ms pre-sound time) or prior to sound onset for movement signals (100ms prior to sound time vs 400-500ms prior to sound time). (D) Functional characterization of select cluster groups. From left to right: PSTHs of average neural response to expected (cyan) and probe frequency (orange) sounds in the active (light) and passive (dark) conditions, and on omission trials (gray). Note different Y axis scales. Quantification of average response strength across all neurons to each sound type and movement signals as measured in (C). Quantification of modulation of each neuron to expected and probe frequency sounds, with neurons included if they have a significant response to the given tone in either the active or passive condition. Right, quantification of group neuron counts by depth (top, black) and fraction of all neurons in each layer (bottom, blue). Note, L4 is omitted due to lack of neurons.
